# Eco-evolutionary dynamics of sexually selected infanticide facilitate species coexistence

**DOI:** 10.64898/2026.10.28.762487

**Authors:** Tatsumi Kudo, Kazuya Kobayashi

## Abstract

Understanding how biodiversity is maintained is a central question in community ecology. Modern coexistence theory predicts that stable coexistence occurs when stabilizing mechanisms are strong enough to overcome competitive ability differences. Recent studies suggest that adaptations to conspecific interactions may provide such a stabilizing mechanism when their demographic costs increase with conspecific density. We examined whether sexually selected male infanticide can promote species coexistence by generating density-dependent demographic costs. We developed a stage-structured eco-evolutionary model of two species competing for a shared resource, with infanticidal and non-infanticidal male strategies, and confirmed that the infanticidal strategy was maintained depending on the parameters related to intraspecific competition: the infanticide rate and its fecundity cost. In a competitively superior species, the infanticidal strategy, which imposed demographic costs that increase with conspecific density, was maintained at a sufficiently high frequency, thereby strengthening self-limitation and reducing the species’ equilibrium density. This reduction increased the invasion growth rate of the inferior competitor and expanded the parameter region permitting stable coexistence. These results show that sexually selected infanticide favored at the individual level can generate population-level self-regulation and alter community-level coexistence.

## Introduction

Understanding the mechanisms that maintain biodiversity is a central question in ecology [1–3]. Although competitive exclusion is expected among species that use the same or similar resources, many competing species coexist over long periods in natural communities [3–5]. Modern coexistence theory explains species coexistence in terms of stabilizing mechanisms, which generate a rare-species advantage, and equalizing mechanisms, which reduce average fitness differences among species [6,7]. When conspecific density has a stronger negative effect on the per-capita population growth than heterospecific density, a species experiences stronger inhibition as its population size increases, resulting in negative frequency dependence at the community level [8,9]. To date, mechanisms underlying the negative frequency dependence have traditionally been attributed to resource partitioning, species-specific predation, and spatiotemporal environmental variation [10–12].

However, the self-limitation underlying such stabilizing effects need not arise solely from external factors. It may also emerge as a demographic consequence of traits shaped by intraspecific interactions [13–15]. In the field of behavioral ecology, which mainly deals with the intraspecific interactions, it is known that certain traits and behaviors increase the relative fitness of the individuals expressing them while causing fitness losses in the other individuals, thereby suppressing the population growth [16–18]. Conflicts between individuals and the resulting population-level performance have been studied within the contexts of the tragedy of the commons in evolutionary biology [19–21]. This demographic load imposed by intraspecific conflicts is proportional to the frequency of conspecific interactions. Consequently, this load increases with conspecific density, resulting in negative density dependence on the per-capita population growth. When the population size of a given species is low, the frequency of conspecific interaction decreases, thereby alleviating this load. Because this intraspecific adaptation load strengthens population self-regulation independently of interspecific competition, it can act as a stabilizing effect that promotes coexistence [6,13,22].

Among such adaptations to conspecific interactions, sexual selection provides a particularly relevant case because they can increase individual reproductive success while simultaneously imposing fitness costs on conspecifics through male mating harassment and disruption of parental care [18,19,22]. When their demographic costs increase with conspecific density, sexually selected traits may generate negative density dependence in population growth and thereby strengthen population self-regulation [23]. However, it remains poorly understood whether behaviors maintained by sexual selection can strengthen population self-regulation and thereby change the invasion growth rates and coexistence of competing species.

Here, we focus on sexually selected male infanticide as an example of this process. Sexually selected infanticide refers to the killing by a male of offspring that he is unlikely to have sired, thereby terminating maternal care, accelerating the female’s return to fertility, and increasing his opportunity to sire her subsequent offspring [24–26]. At the population level, infanticide eliminates ongoing reproductive investment and increases the effective time required to produce the next generation, thereby reducing female lifetime reproductive success and population growth rates [27,28]. If infanticide is mediated by encounters between caring females and infanticidal males, and the frequency of such encounters increases with conspecific density, its demographic cost should be greater at higher densities. Thus, male infanticide provides a tractable example of precisely this conflict between individual-level advantage and population-level demographic cost.

These features of infanticide lead to two questions: under what conditions can infanticide evolve and be maintained by sexual selection, and once maintained, can its demographic cost alter the coexistence of competing species? In this study, we develop a stage-structured eco-evolutionary model of two species competing for a shared resource. Within each species, females transition among a free state and two sequential stages of parental care, whereas males express either an infanticidal or a non-infanticidal strategy. Infanticide occurs through conspecific encounters between caring females and infanticidal males, terminates ongoing parental care, and resets reproduction to the initial care stage with a new brood sired by the infanticidal male. We first analyze the one-species model to identify the conditions under which the infanticidal strategy evolves and to quantify the resulting changes in equilibrium population density. We then analyze the two-species competition model to determine whether infanticide-induced demographic costs generate negative density dependence in population growth and thereby strengthen population self-regulation and affect species coexistence.

## Models

### 1. Life history

We consider two closely related animal species, *i* ∈ {1,2}, that compete for a shared resource, with competition represented by a Lotka–Volterra formulation [2]. In both species, only females provide parental care, and males express one of two strategies: an infanticidal strategy *I*, or a non-infanticidal strategy *N*. We assume that reproduction and infanticide occur only between conspecific individuals, whereas interspecific interactions occur exclusively through resource competition.

The adults of species *i* are divided into five female classes and two male classes (Fig. 1). Females are classified as either free females *F*_*f,i*_ or caring females. Caring females are further classified according to the stage of parental care (first or second) and the strategy of the male that sired their offspring (non-infanticidal or infanticidal), yielding four classes: *C*_1,*N,i*_, *C*_2,*N,i*_, *C*_1,*I,i*_, and *C*_2,*I,i*_. Males are classified as non-infanticidal *M*_*N,i*_ or infanticidal *M*_*I,i*_.

**Figure 1.**
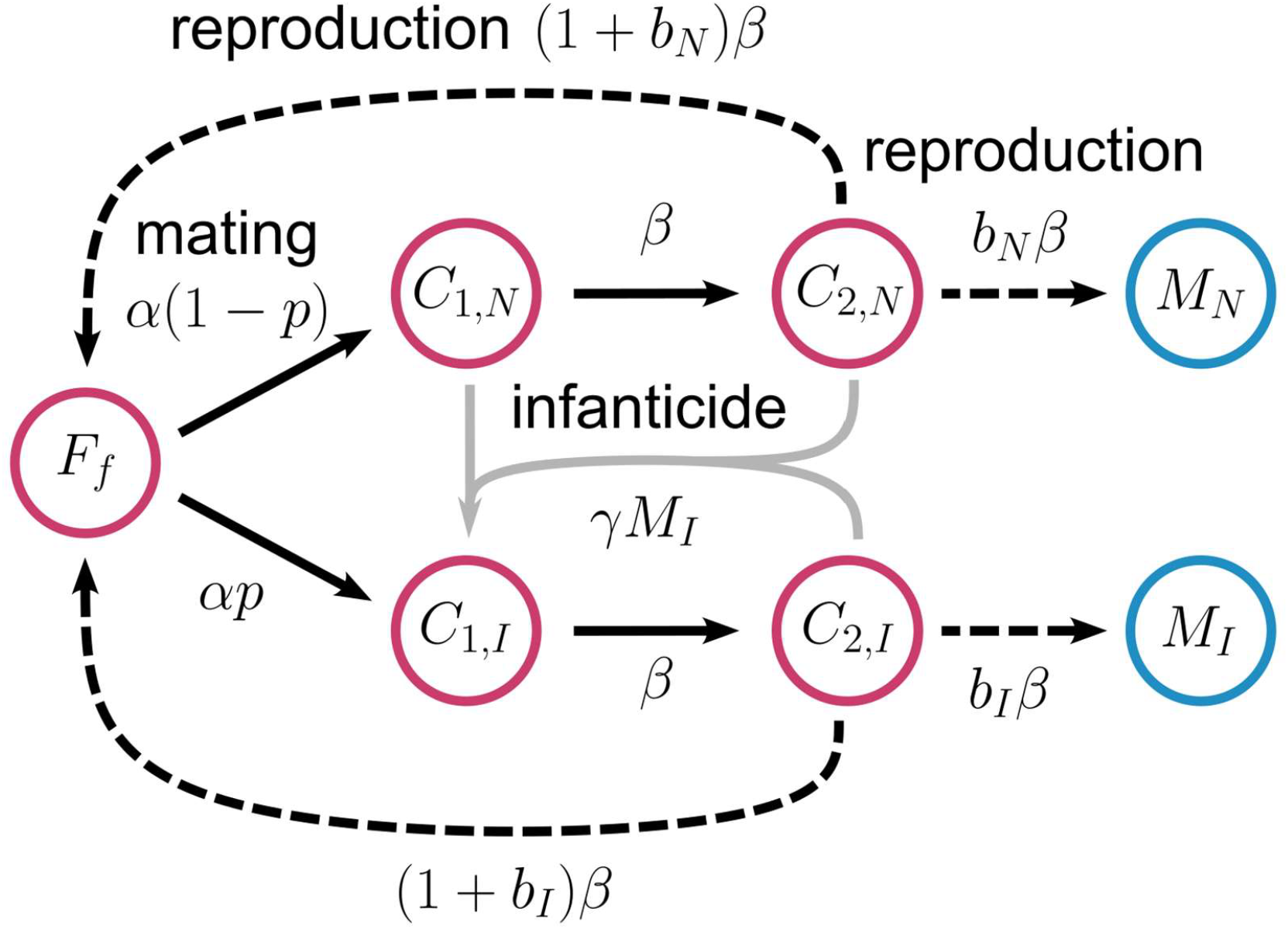
Schematic diagram of the stage-structured model. Red circles indicate female classes, and blue circles indicate male classes. *F*_*f*_ is the density of free females; *C*_1,*N*_ and *C*_2,*N*_ are females in the first and second care stages with broods sired by non-infanticidal males; *C*_1,*I*_ and *C*_2,*I*_ are the corresponding classes with broods sired by infanticidal males; and *M*_*N*_ and *M*_*I*_ are the densities of non-infanticidal and infanticidal males, respectively. Solid arrows indicate stage transitions, and dashed arrows indicate reproduction. All classes experience density-dependent mortality at rate *dN*.

We assume that free females initiate reproduction at the rate *αi* independent of male density and mate randomly with respect to male strategy. The frequency of infanticidal males is defined as *pi* = *M*_*I,i*_/(*M*_*N,i*_ + *M*_*I,i*_). Accordingly, the females mate with an infanticidal male with probability *pi* and with a non-infanticidal male with probability 1 − *p*_*i*_. The treatment of extinction boundaries, at which this frequency is undefined, is described in the Supplementary Materials.

We assume that successful offspring recruitment requires the completion of parental care without interruption by infanticide. If parental care were represented by a single Markovian compartment, returning a female to the same care compartment after an infanticide would not change her expected remaining time to reproductive completion, making the demographic effect of interrupted care invisible. We therefore use the linear chain trick to represent parental care as two sequential stages, allowing infanticide to erase progress accumulated toward successful offspring recruitment [29,30]. Females progress from the first to the second care stage at rate *β* and complete parental care from the second stage at the same rate *β*. Upon successful completion, the mother returns to the free-female class, and adult daughters and sons are recruited into the population.

We assume a primary sex ratio of 1:1. Each completed reproductive event sired by a male of strategy *k* ∈ {*N, I*} produces *b*_*k,i*_ adult daughters and *b*_*k,i*_ adult sons. Sons are assumed to inherit their father’s strategy, corresponding to complete Y-linked inheritance of the male strategy.

Infanticide occurs when a caring female encounters a conspecific infanticidal male. Assuming mass-action mixing, each caring female experiences a density-dependent infanticide rate of *γ*_*i*_*M*_*I,i*_. When infanticide occurs, the current brood is lost and replaced by a new brood sired by the infanticidal male, and the female enters the first care stage, *C*_1,*I,i*_. In the Supplementary Materials, we additionally analyze alternative models in which reproductive initiation is also density dependent or infanticide is density independent. We further assume that the infanticidal strategy incurs a fecundity cost. The reproductive output of broods sired by infanticidal males is given by *b*_*I,i*_ = (1 − *s*_*i*_)*b*_*N,i*_, where 0 v *s*_*i*_ v 1 represents the proportional reduction in offspring production relative to broods sired by non-infanticidal males.

Based on the life history described above, the eco-evolutionary dynamics of species *i* are written as

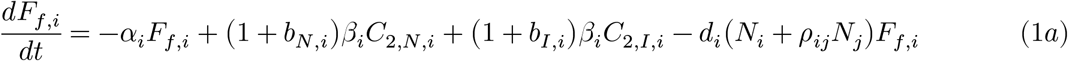

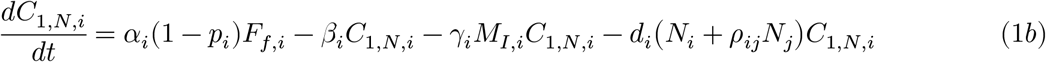

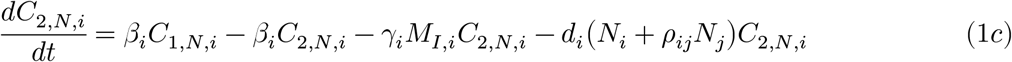

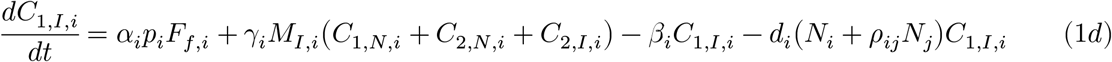

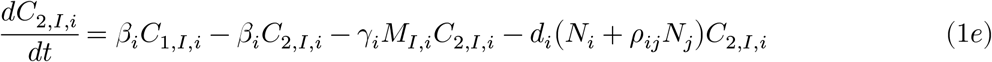

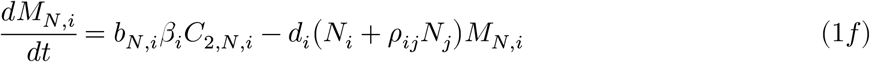

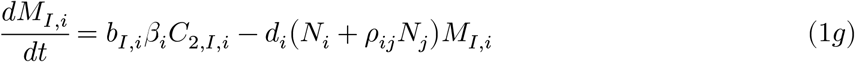

where *N*_*i*_ is the total adult density of species *i, d*_*i*_ is the strength of density-dependent mortality and *ρ*_*ij*_ represents the competitive effect of species *j* on species *i*.

### 2. Analysis

We numerically identified biologically feasible equilibria of the one- and two-species systems, as closed-form solutions could not be obtained for all equilibria. Local stability was assessed using the eigenvalues of the corresponding Jacobian matrices. Equilibria were classified as locally asymptotically stable when all eigenvalues had negative real parts and as unstable when at least one eigenvalue had a positive real part [31].

For the one-species system, we set the competing species’ density to zero and examined the equilibria and invasion thresholds of the two male strategies. The invasion thresholds were derived analytically through evolutionary invasion analysis, using the dominant eigenvalues of linearized rare-strategy subsystems [32–35]. Analytical expressions for boundary equilibria and invasion conditions are provided in the Supplementary Materials.

For the two-species system, we first fixed male-strategy frequencies and applied a quasi-steady-state approximation based on a separation of time scales to reduce the stage-structured dynamics to a two-dimensional system of total population densities. We assumed that the adult sex ratio is fixed at 1:1 and that the distribution of females among reproductive stages reaches a quasi-steady state much faster than the total population density changes (Eq. S37 in Supplementary Materials). We used this reduced model to analyze nullclines, equilibria, and phase-plane dynamics. We also assessed mutual invasibility in Supplementary Materials. We then analyzed the full 14-dimensional eco-evolutionary model to examine how infanticide evolution affects species coexistence. We assessed the extent to which the evolution of infanticide expanded the parameter region permitting species coexistence by comparing this region with that of a model without the infanticidal strategy. At species-extinction equilibria, stability was evaluated using the resident-species Jacobian and the invasion growth rate of the missing species, because male-strategy frequencies are undefined at extinction. Species differed only in reproductive output, with *b*_*N*,1_ > *b*_*N*,2_.

All numerical analyses were conducted in Python using NumPy and SciPy [36,37]. Representative trajectories were obtained by numerically integrating the ordinary differential equations. Each parameter range was divided into 100 intervals, and equilibrium searches and stability analyses were repeated to construct bifurcation diagrams and parameter planes (Table 1). Parameter planes were classified by the types of locally asymptotically stable equilibria. Parameter values and ranges are provided in the corresponding figure captions. The code required to reproduce all analyses and figures is available from the GitHub repository (https://github.com/putaro-cu/EcoEvoInfanticide).

**Table 1.** Model variables and parameters. The parameters within brackets were varied over their respective analyzed ranges, each divided into 100 equal intervals, while all the others were kept at the specified values.

| Symbol | Definition | Analyzed range |
| --- | --- | --- |
| $F_{f,i}$ | Density of free females of species $i$ | |
| $C_{1,N,i}$ | Density of females of species $i$ in the first parental-care stage, caring for offspring sired by a non-infanticidal male | |
| $C_{2,N,i}$ | Density of females of species $i$ in the second parental-care stage, caring for offspring sired by a non-infanticidal male | |
| $C_{1,I,i}$ | Density of females of species $i$ in the first parental-care stage, caring for offspring sired by an infanticidal male | |
| $C_{2,I,i}$ | Density of females of species $i$ in the second parental-care stage, caring for offspring sired by an infanticidal male | |
| $M_{N,i}$ | Density of non-infanticidal males of species $i$ | |
| $M_{I,i}$ | Density of infanticidal males of species $i$ | |
| $N_i$ | Total adult density of species $i$ | |
| $p_i$ | Frequency of infanticidal males in species $i$ | |
| $\alpha_i$ | Per-capita rate at which a free female of species $i$ initiates reproduction | 1 |
| $\beta_i$ | Transition rate between consecutive parental-care stages and the rate of completion of the second care stage | 1 |
| $\gamma_i$ | Mass-action coefficient determining the rate of infanticidal encounters in species $i$ | [0,0.02] |
| $b_{N,i}$ | Number of adult daughters (equal to the number of adult sons) recruited from a brood sired by a non-infanticidal male ($b_{N,1} > b_{N,2}$) | [0,2] |
| $b_{I,i}$ | Number of adult daughters (equal to the number of adult sons) recruited from a brood sired by an infanticidal male | [0,2] |
| $s_i$ | Proportional reduction in the reproductive output of broods sired by infanticidal males | [0,1] |
| $d_i$ | Coefficient of density-dependent mortality in species $i$ | 0.001 |
| $\rho_{i,j}$ | Competition coefficient of species $j$ on species $i$ | [0,1] |

## Results

### 1. one-species model

Across the parameter ranges examined, there were up to four biologically feasible equilibria: the extinction equilibrium *E*_0_, two boundary equilibria *E*_*N*_ and *E*_*I*_, corresponding to fixation of the non-infanticidal and infanticidal strategies, respectively, and at most one interior equilibrium *E*_*P*_. Stability analysis revealed that the extinction equilibrium *E*_0_ was always unstable, whereas exactly one of the remaining equilibria, *E*_*N*_, *E*_*I*_, or *E*_*P*_, was locally stable for each parameter combination examined (Fig. 2). Accordingly, the trajectories converged to *E*_*N*_, *E*_*I*_, or *E*_*P*_, depending on the parameter values (Fig. 3).

**Figure 2.**
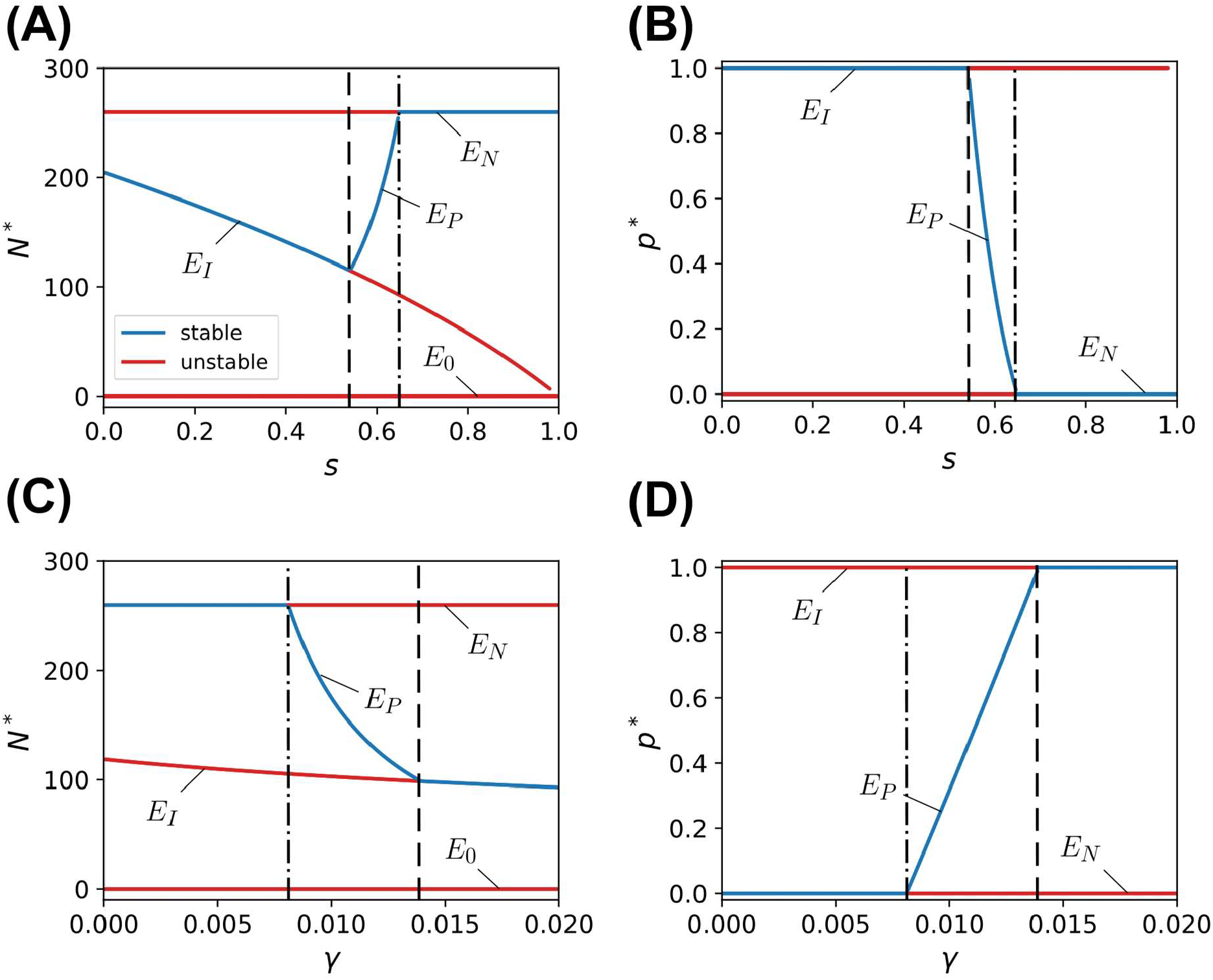
Bifurcation diagrams of demographic and evolutionary equilibria with respect to the cost *s* and the rate *γ* of infanticide strategy. Equilibrium frequency of infanticidal males, *p*^∗^, and total population density, *N*^∗^, are plotted against the parameters. (A, B) show bifurcations along *s*, the reduction in fecundity of infanticidal males. (C, D) show bifurcations along ***γ***, the rate of infanticide. Blue and red branches indicate local stability: stable and unstable equilibria, respectively. The dash-dotted and dashed lines indicate the invasion criteria in Eq. 2 and 3, respectively. In (A, B), ***γ*** = 0.01, and in (C, D), *s* = 0.6. Other parameters are *α* = 1, *β* = 1, *d* = 0.001, *b*_*N*_ = 1.

**Figure 3.**
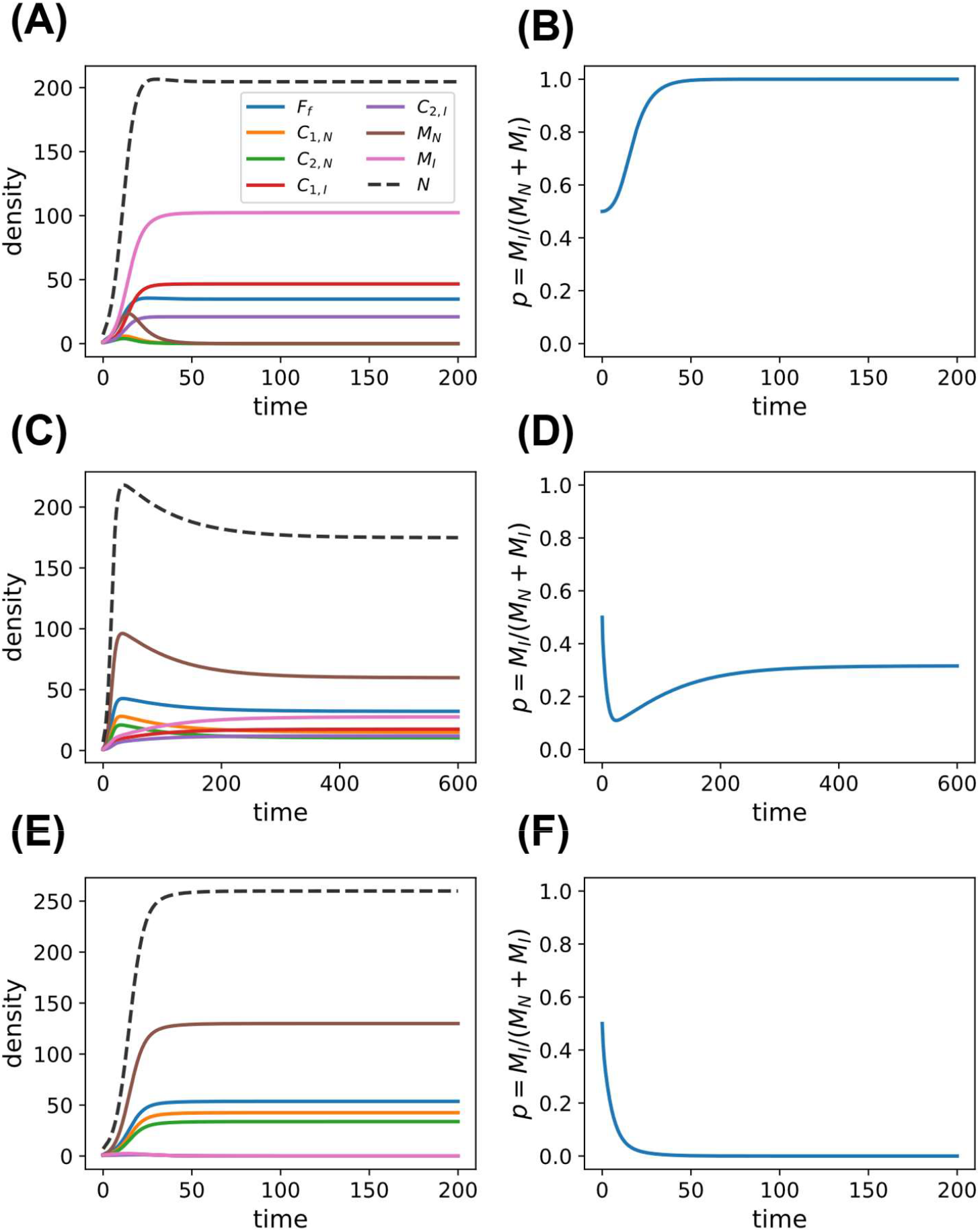
Eco-evolutionary dynamics of the one-species model under different costs of the infanticidal strategy. (A, C, E) show the population dynamics of all state variables and the total density *N*, and (B, D, F) show the temporal change in the frequency of infanticidal males, *p* = *M*_*I*_/(*M*_*N*_ + *M*_*I*_). (A, B) In the absence of cost, *s* = 0, the infanticidal strategy is fixed in the population (*p*^∗^ = 1). (C, D) At intermediate cost, *s* = 0.6, both strategies are maintained, yielding a stable polymorphic equilibrium (0 < *p*^∗^ < 1). (E, F) At high cost, *s* = 0.8, the non-infanticidal strategy is fixed (*p*^∗^ = 0). Other parameters are *α* = 1, *β* = 1, *γ* = 0.01, *d* = 0.001, and *b*_*N*_ = 1.

We performed an evolutionary invasion analysis, in which the invasion fitness of a rare mutant is given by the dominant eigenvalue of the linearized mutant subsystem in the environment established by the resident population. Detailed derivations of these invasion criteria are provided in the supplementary material. At the non-infanticidal boundary equilibrium *E*_*N*_, the infanticidal strategy could invade when

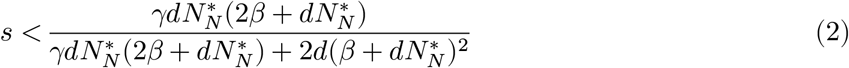

where *s* is the fecundity cost of the infanticidal strategy, *γ* is the infanticide rate, and 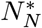 is the total population density at the non-infanticidal boundary equilibrium and is the root of the cubic equation (Eq. S4).

Conversely, at the infanticidal boundary equilibrium *E*_*I*_, the non-infanticidal strategy could invade when

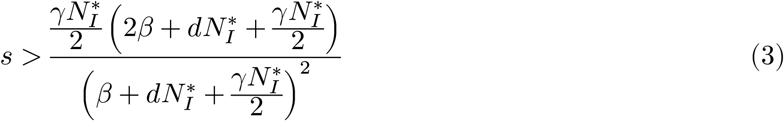

Here, 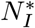 is the total population density at the infanticidal boundary equilibrium and is the root of the cubic equation (Eq. S5). Note that 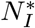 depends implicitly on *s*.

We identified the equilibria numerically and evaluated their local stability from the eigenvalues of the corresponding Jacobian matrices. Consistent with the invasion criteria in Eq. 2 and Eq. 3, three evolutionary outcomes were observed (Fig. 2). When only the infanticidal strategy could invade the alternative boundary equilibrium, the infanticidal strategy became fixed. When only the non-infanticidal strategy could invade, the non-infanticidal strategy became fixed. When both strategies could invade each other’s boundary equilibrium, a locally stable internal polymorphic equilibrium was established (Fig. 2).

### 2. two-species model

We first analyzed a non-evolutionary model in which the frequency of infanticidal males, *p*_*i*_, was held fixed in each species to separate the ecological effects of infanticide on interspecific competition from changes in male-strategy frequencies. We reduced the original stage-structured model to a two-dimensional system for the total population densities *N*_1_ and *N*_2_ using a quasi-steady state approximation. Detailed derivations are provided in the Supplementary Materials.

Figure 4 shows the ecological outcomes of the reduced system as the frequencies of infanticidal males in the two species, *p*_1_ and *p*_2_, were varied. Across the parameter range examined, the system exhibited either stable coexistence or competitive exclusion of the competitively inferior species 2. When the frequency of infanticidal males in the superior species 1, *p*_1_, was low, species 2 was competitively excluded (Figs. 4B, C), whereas increasing *p*_1_ led to the emergence of a locally stable coexistence equilibrium and extended the parameter region permitting coexistence (Figs. 4D, E). In contrast, increasing *p*_2_ reduced the region permitting coexistence. Thus, the frequencies of infanticidal males in the two species had opposing effects on coexistence: infanticide in the competitively superior species promoted coexistence, whereas infanticide in the inferior species restricted it.

**Figure 4.**
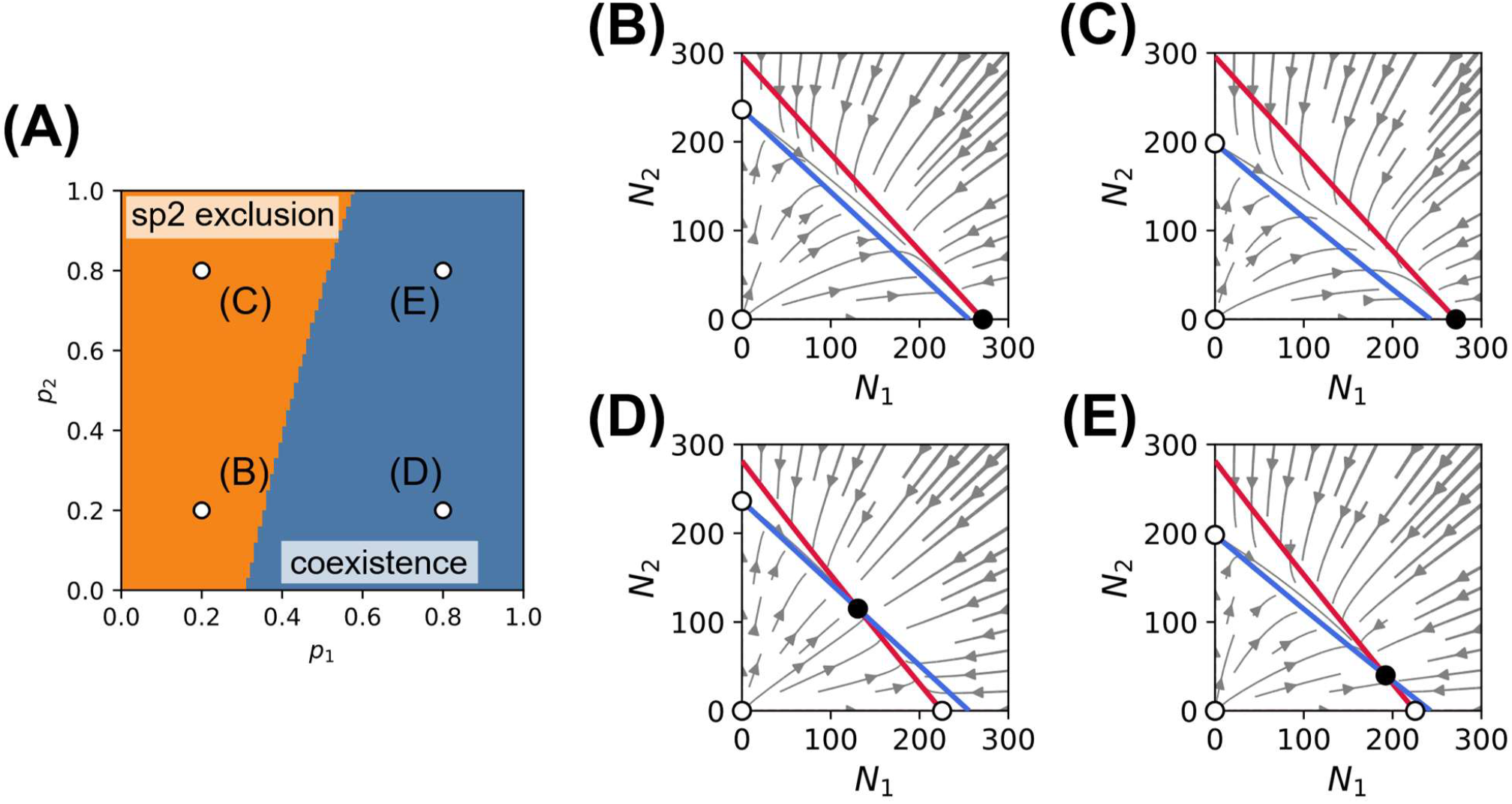
Ecological outcomes and phase planes of the reduced two-species population dynamics under fixed infanticidal male frequencies. Population dynamics were reduced to a two-dimensional system for the total population densities *N*_1_ and *N*_2_ by a time-scale separation approximation. (A) Outcome plane as a function of the fixed frequencies of infanticidal males in species 1 and species 2, *p*_1_ and *p*_2_, respectively. Blue indicates locally stable coexistence, whereas orange indicates competitive exclusion of species 2. Open circles in (A) indicate the parameter combinations shown in the phase portraits: (B) (*p*_1_, *p*_2_) = (0.2, 0.2), (C) (*p*_1_, *p*_2_) = (0.2, 0.8), (D) (*p*_1_, *p*_2_) = (0.8, 0.2), and (E) (*p*_1_, *p*_2_) = (0.8, 0.8). In panels (B–E), streamlines indicate the direction of population change in the *N*_1_ –*N*_2_ phase plane. Red and blue curves denote the non-trivial nullclines *dN*_1_/*dt* = 0 and *dN*_2_/*dt* = 0, respectively. Filled circles indicate locally stable equilibria, whereas open circles indicate unstable equilibria. Other parameters are *b*_*N*,1_ = 1.2, *b*_*N*,2_ = 1.0, *α*_1_ = *α*_2_ = 1, *s* = 0.1, *β*_1_ = *β*_2_= 1, *γ*_1_ = *γ*_2_ = 0.01, *d*_1_ = *d*_2_ = 0.001, and *ρ*_12_ = *ρ*_21_ = 1.

We next examined whether the evolution of infanticidal strategies could promote species coexistence using the full two-species eco-evolutionary model, in which male-strategy frequencies evolve within each species. Our numerical analysis indicated that the system had a unique locally stable equilibrium for each parameter combination within the parameter ranges examined. We then classified the ecological and evolutionary outcomes based on the type of this locally stable equilibrium (Fig. 5).

**Figure 5.**
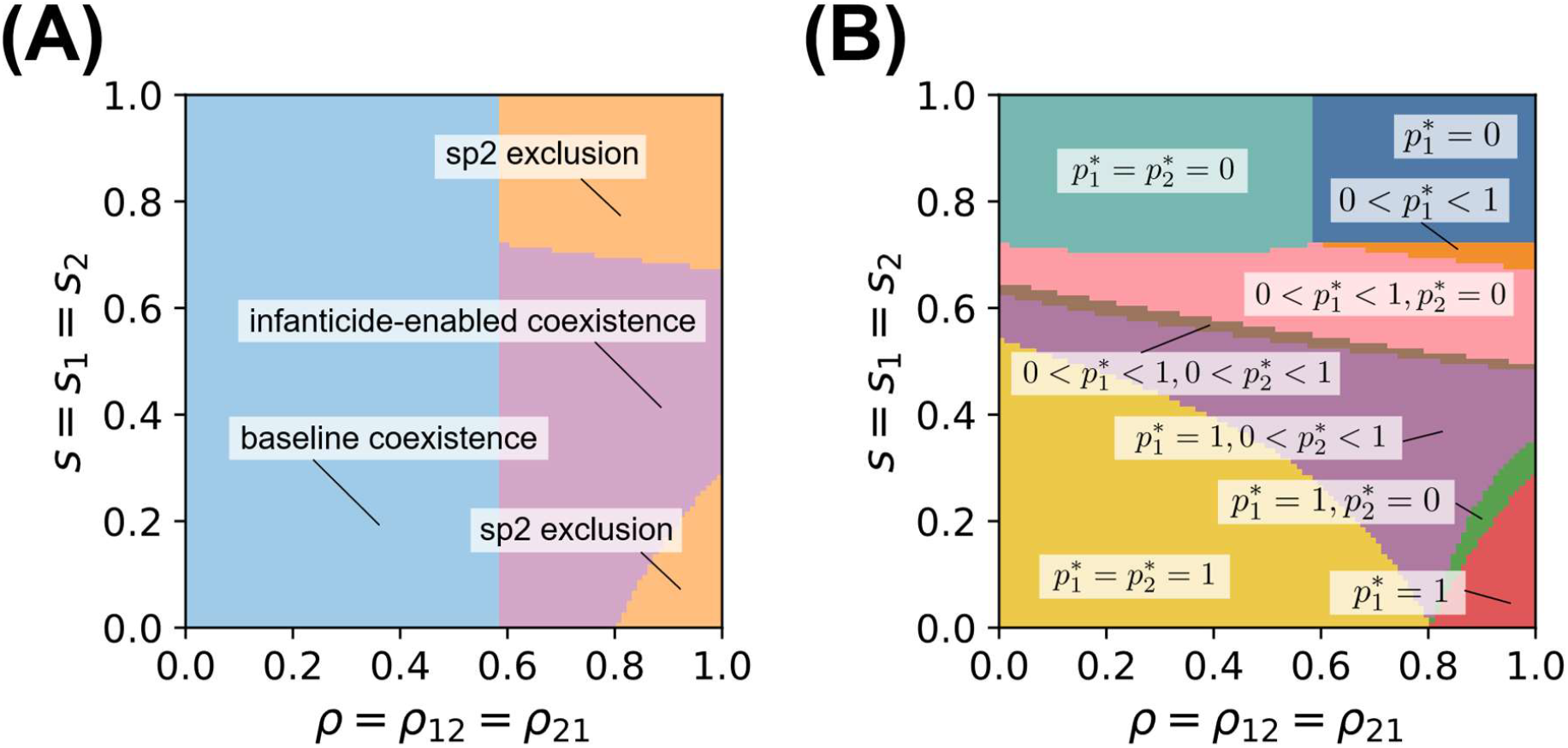
Infanticide-driven expansion of species coexistence and eco-evolutionary outcomes in the full two-species model. Classification of locally stable equilibria in the parameter plane defined by the interspecific competition coefficient, *ρ* = *ρ*_12_ = *ρ*_21_, and the fecundity cost of the infanticidal strategy, *s* = *s*_1_ = *s*_2_. Only one stable equilibrium was identified for each parameter combination examined. (A) Colors indicate ecological outcomes. Baseline coexistence refers to the region where a stable coexistence equilibrium exists even when both species consist entirely of non-infanticidal individuals. The evolution of infanticide expanded the parameter region in which a stable coexistence equilibrium exists (purple region). (B) Colors indicate the equilibrium frequencies of infanticidal males in the stable equilibrium. In the red, blue, and orange regions, 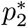 is undefined because species 2 is extinct. Other parameters are *b*_*N*,1_ = 2.0, *b*_*N*,2_ = 1.0, *α*_1_ = *α*_2_ = 1, *β*_1_ = *β*_2_ = 1, *γ*_1_ = *γ*_2_ = 0.01, and *d*_1_ = *d*_2_ = 0.001.

The ecological outcomes were classified into two categories: coexistence of the two species and competitive exclusion of species 2 (Fig. 5A). The coexistence region was further divided into baseline coexistence, where the two species could coexist even in the absence of infanticide, and infanticide-enabled coexistence, where coexistence was possible due to the presence of infanticide. Infanticide-enabled coexistence occurred at low to intermediate fecundity costs *s*, and the range of *s* permitting coexistence narrowed as the interspecific competition coefficient *ρ* increased. Notably, at sufficiently high *ρ*, species 2 was competitively excluded at low *s*, whereas stable coexistence was possible at intermediate *s*.

The corresponding evolutionary outcomes also varied across the parameters (Fig. 5B). At low fecundity costs *s*, the infanticidal strategy became fixed in both species. As *s* increased, the evolutionary outcomes shifted from fixation of the infanticidal strategy to stable polymorphism and eventually to fixation of the non-infanticidal strategy, as observed in the one-species model. However, at high *ρ*, a different pattern was observed in species 2: as *s* increased, the non-infanticidal strategy became fixed, followed by stable polymorphism, and then fixation of the non-infanticidal strategy again (*ρ* > 0.8 in Fig. 5B).

Comparing the ecological and evolutionary outcomes revealed that the infanticide-enabled coexistence region was associated with the maintenance of the infanticidal strategy in species 1 (Fig. 5). When the infanticidal strategy was maintained at a sufficiently high frequency in the competitively superior species, stable coexistence was allowed under interspecific competition conditions that would otherwise lead to the exclusion of species 2 in the absence of infanticide. In contrast, at the high fecundity cost, the frequency of infanticidal males in species 1 declined, and species 2 was competitively excluded. Furthermore, even when the infanticidal strategy was fixed in species 1, species 2 was competitively excluded if the interspecific competition coefficient *ρ* was sufficiently high (orange region in the lower-right corner in Fig. 5A).

## Discussion

In this study, we showed that the eco-evolutionary dynamics of sexually selected male infanticide can generate an additional source of negative density dependence in per capita population growth, thereby strengthening population self-regulation and expanding the parameter region in which competing species can coexist stably. Three main results support this conclusion. First, the infanticidal strategy could evolve through its reproductive advantage at the individual level, while its evolution reduced equilibrium population density (Figs. 2 and 3). Second, when infanticidal males were present at sufficiently high frequency in the competitively superior species, the reduction in its equilibrium population density facilitated the invasion of the competitively inferior species and enabled stable coexistence (Fig. 4). Third, even when both population densities and male-strategy frequencies were allowed to change dynamically in the full eco-evolutionary model, the infanticidal strategy was maintained and stable species coexistence was achieved (Fig. 5).

The stable polymorphism observed in the one-species model can be explained by negative frequency-dependent selection mediated by population density. When infanticidal males become more frequent, interruptions of maternal care reduce population density, thereby decreasing opportunities for paternity acquisition through infanticide and favoring the non-infanticidal strategy, which bears no fecundity cost. Conversely, when infanticidal males become less frequent, population density increases, enhancing the reproductive advantage of infanticide. Thus, at intermediate fecundity costs, eco-evolutionary feedback between strategy frequency and population density can maintain a stable polymorphism of infanticide behavior.

Infanticide imposed demographic costs at the population level not only through the fecundity cost associated with the strategy itself but also by interrupting ongoing maternal care and delaying reproductive completion. Our model captures the temporal component of such lost reproductive investment using the linear-chain trick, which allows an infanticide event to erase progress already made through the parental-care process and return the female to an earlier reproductive stage. Consequently, infanticide reduces the number of successful reproductive events completed per unit time. Indeed, equilibrium population density declined even when the two male strategies had identical reproductive outputs, indicating that reproductive delay alone can suppress population growth (*E*_*N*_ and *E*_*I*_ branches in Fig. 2A, Eq. S7 in Supplementary Materials). Such reproductive losses are consistent with empirical observations in white-faced capuchins, where infant loss imposes substantial temporal and energetic costs, and females exposed to more frequent male replacements associated with elevated infanticide risk tend to produce fewer surviving offspring over their lifetimes [38].

The demographic effects of infanticide influenced species coexistence by strengthening intraspecific self-limitation relative to interspecific competition. In the reduced model with fixed male-strategy frequencies, the rate of infanticide increased with conspecific density, causing more frequent interruptions of maternal care and reducing the effective recruitment rate. Because this additional demographic cost weakened as a species became rare, density-dependent infanticide could generate a rare-species advantage, consistent with the stabilizing mechanisms described by modern coexistence theory [6,7]. Infanticide in the competitively superior species reduced its single-species equilibrium density, relieving interspecific competition and increasing the invasion growth rate of the inferior species. In the simplified case without a direct fecundity cost, invasion analysis further showed that this reduction in resident density could allow the inferior species to invade, promoting mutual invasibility (Supplementary Materials). Consistent with this mechanism, the reduced model exhibited an expanded region of stable coexistence (Fig. 4). Importantly, the full eco-evolutionary model further showed that sexual selection could maintain infanticidal males at frequencies sufficient to promote stable coexistence, even when male-strategy frequencies and population densities changed dynamically (Fig. 5). These results suggest that sexually selected infanticide can act as a stabilizing mechanism promoting species coexistence by generating density-dependent demographic costs that strengthen intraspecific self-limitation.

The stabilizing mechanism identified in our model relies on the simple assumption that the rate of infanticide increases linearly with conspecific population density. The relationship between infanticide and population density in natural populations may be influenced by territoriality, spatial structure, mating systems, and other ecological factors [39–41]. However, empirical studies have documented density-dependent increases in infanticide, suggesting that this assumption is biologically plausible, although the strength and functional form of this relationship may vary among species [41–43]. Furthermore, our model focuses on sexually selected infanticide and does not consider other potential benefits of infanticide or the evolution of female counterstrategies. Future studies should examine whether density-dependent infanticide can promote species coexistence under more realistic ecological and evolutionary conditions.

More generally, our findings suggest that sexually selected infanticide represents a form of intraspecific adaptation load, in which a behavior favored by individual-level reproductive benefits imposes demographic costs at the population level [13]. Unlike mechanisms involving resource partitioning or changes in interspecific competition coefficients, infanticide strengthened intraspecific self-limitation through sexual interactions. Our results suggest that other demographically costly intraspecific interactions, such as cannibalism, territorial competition, and intraspecific aggression, may also contribute to the stabilizing mechanisms underlying species coexistence when their demographic costs increase with conspecific density [44–47].

Testing this mechanism in natural communities will require empirical systems in which reproductive interactions, their demographic consequences, and population density can be quantified or experimentally manipulated. Burying beetles (*Nicrophorus spp*.) and some mammals, in which infanticide has been documented and which coexist with competitors exploiting similar resources, may provide promising systems for such tests [48,49]. Future studies should examine whether density-dependent reproductive costs associated with infanticide generate sufficient self-limitation to facilitate the invasion of competing species. More broadly, theoretical and empirical work is needed to determine whether adaptations to conspecific interactions other than infanticide similarly contribute to species coexistence in natural communities.

## Data accessibility

All source codes used in this article are available in the GitHub repository at https://github.com/putaro-cu/EcoEvoInfanticide.

## Declaration of AI use

Artificial intelligence was used to improve the quality of the code and English writing in this article.

## Acknowledgements

We are grateful to S. Dobata for valuable comments and suggestions.

## Funding statement

This research was partly supported by Grant-in-Aid for JSPS Fellows No. 26KJ0192 to T.K.

## Competing interests

The authors declare no competing interests.

## Notes

https://github.com/putaro-cu/EcoEvoInfanticide

### Competing Interest Statement

The authors have declared no competing interest.

