## Supplementary Materials for "Eco-evolutionary dynamics of sexually selected infanticide facilitate species coexistence"

### 1. one-species model

In the one-species model, there are three boundary equilibria: the extinction equilibrium and two monomorphic equilibria corresponding to $p = 0$ and $p = 1$. We denote these equilibria by $E_0$, $E_N$, and $E_I$, respectively.

#### 1.1 extinction equilibria

At the extinction equilibrium $E_0$, the male-strategy frequency $p = M_I/(M_N + M_I)$ is undefined because $M_N = M_I = 0$. Although the vector field can be continuously extended to $E_0$ by setting $\boldsymbol{f}(E_0) = \mathbf{0}$, the extended vector field is not differentiable at the origin. Hence, a unique Jacobian of the full system does not exist at $E_0$. To establish instability, however, it is sufficient to consider the invariant non-infanticidal boundary subspace, in which all infanticidal classes remain zero and $p = 0$. Linearization within this subspace gives

$$\lambda[(\lambda+\alpha)(\lambda+\beta)^2 - (1+b_N)\alpha\beta^2] = 0. \tag{S1}$$

Let $Q(\lambda) = (\lambda+\alpha)(\lambda+\beta)^2 - (1+b_N)\alpha\beta^2$. For $b_N > 0$, $Q(0) = -b_N\alpha\beta^2 < 0$, whereas $Q(\lambda) \to \infty$ as $\lambda \to \infty$. Because $Q'(\lambda) > 0$ for $\lambda \geq 0$, $Q$ has a unique positive root. Thus, the restricted system has a positive eigenvalue and therefore a biologically feasible direction along which trajectories move away from extinction. Hence, $E_0$ is unstable in the full system.

#### 1.2 boundary equilibria

The non-infanticidal boundary equilibrium $E_N = (F_N^*, C_{1,N}^*, C_{2,N}^*, 0{,}0, M_N^*, 0)$ and the infanticidal boundary equilibrium $E_I = (F_I^*, 0{,}0, C_{1,I}^*, C_{2,I}^*, 0, M_I^*)$, can be obtained by solving Eq. 1:

$$E_N = \left(\frac{d(1+b_N)(N_N^*)^2}{2b_N(\alpha+dN_N^*)}, \frac{d(N_N^*)^2(\beta+dN_N^*)}{2b_N\beta^2}, \frac{d(N_N^*)^2}{2b_N\beta}, 0{,}0, \frac{N_N^*}{2}, 0\right), \tag{S2}$$

and

$$E_I = \left(\frac{d(1+b_I)(N_I^*)^2}{2b_I(\alpha+dN_I^*)}, 0{,}0, \frac{d(N_I^*)^2(\beta+dN_I^*+\gamma N_I^*/2)}{2b_I\beta^2}, \frac{d(N_I^*)^2}{2b_I\beta}, 0, \frac{N_I^*}{2}\right). \tag{S3}$$

Here, $N_N^*$ and $N_I^*$ denote the equilibrium total population densities at the non-infanticidal and infanticidal boundary equilibria, respectively. They are determined by

$$d^3(N_N^*)^3 + d^2(\alpha + 2\beta)(N_N^*)^2 + d\beta(2\alpha + \beta)N_N^* - \alpha\beta^2 b_N = 0, \tag{S4}$$

and

$$d^2\left(d + \frac{\gamma}{2}\right)(N_I^*)^3 + d\left[d(\alpha + 2\beta) + \frac{\alpha\gamma}{2}\right](N_I^*)^2 + d\beta(2\alpha + \beta)N_I^* - \alpha\beta^2 b_I = 0. \tag{S5}$$

For strictly positive parameter values, each equation admits a unique positive root.

From Eq. S4 and Eq. S5, $b_I$ and $b_N$ satisfy the following relation,

$$\frac{(\alpha + dN_I^*)\left[(\beta + dN_I^*)^2 + \frac{d}{2}\gamma(N_I^*)^2\right]}{1 + b_I} = \frac{(\alpha + dN_N^*)(\beta + dN_N^*)^2}{1 + b_N} = \alpha\beta^2. \tag{S6}$$

Because $b_I \leq b_N$ and $\gamma \geq 0$, the left-hand side for the infanticidal equilibrium is greater than or equal to that for the non-infanticidal equilibrium at any $N > 0$. Since both functions are increasing in $N$,

$$N_I^* \leq N_N^*, \tag{S7}$$

with strict inequality if $b_I < b_N$ or $\gamma > 0$.

#### 1.3 rare-strategy invasion analysis

To characterize the evolutionary outcomes of the one-species model, we performed an evolutionary invasion analysis, in which the invasion fitness of a rare mutant is given by the dominant eigenvalue of the linearized mutant subsystem in the environment established by the resident population [1–4]. A positive invasion fitness indicates that the mutant can invade the resident boundary equilibrium, whereas a negative value indicates resistance to invasion. If only one strategy can invade, that strategy is expected to become fixed; if both strategies can invade, reciprocal invasibility supports a protected polymorphism.

##### 1.3.1 Invasion of the infanticidal strategy into $E_N$

When an infanticidal mutant is rare in a resident population at the non-infanticidal boundary equilibrium $E_N$, the mutant variables are

$$\mathbf{y}_I = (C_{1,I}, C_{2,I}, M_I)^\top. \tag{S8}$$

The linearized mutant subsystem around $E_N$ is

$$\frac{d\mathbf{y}_I}{dt} = A_{I|N}\mathbf{y}_I, \tag{S9}$$

where

$$A_{I|N} = \begin{pmatrix} -(\beta + dN_N^*) & 0 & \dfrac{\alpha F_{f,N}^*}{M_N^*} + \gamma(C_{1,N}^* + C_{2,N}^*) \\ \beta & -(\beta + dN_N^*) & 0 \\ 0 & b_I\beta & -dN_N^* \end{pmatrix}. \tag{S10}$$

Thus, the invasion fitness of the infanticidal mutant is the eigenvalue of $A_{I|N}$ with the largest real part:

$$s_{I|N} = \max_j \operatorname{Re} \lambda_j \left(A_{I|N}\right). \tag{S11}$$

The characteristic polynomial of $A_{I|N}$ is

$$P_{I|N}(\lambda) = (\lambda + \beta + dN_N^*)^2(\lambda + dN_N^*) - b_I\beta^2 \left[\frac{\alpha F_{f,N}^*}{M_N^*} + \gamma(C_{1,N}^* + C_{2,N}^*)\right]. \tag{S12}$$

For $b_I > 0$, $A_{I|N}$ is an irreducible Metzler matrix, and its dominant eigenvalue is real [5]. Because $P_{I|N}(\lambda)$ is strictly increasing for $\lambda \geq 0$, the infanticidal strategy can invade if and only if $P_{I|N}(0) < 0$. Thus,

$$s_{I|N} > 0 \;\Leftrightarrow\; b_I\beta^2 \left[\frac{\alpha F_{f,N}^*}{M_N^*} + \gamma(C_{1,N}^* + C_{2,N}^*)\right] > dN_N^*(\beta + dN_N^*)^2. \tag{S13}$$

Using the equilibrium relations at $E_N$ and $b_I = (1-s)b_N$, this condition reduces to

$$(1-s)\gamma dN_N^*(2\beta + dN_N^*) > 2ds(\beta + dN_N^*)^2 \tag{S14}$$

Equivalently, the infanticidal strategy can invade when

$$s < \frac{\gamma dN_N^*(2\beta + dN_N^*)}{\gamma dN_N^*(2\beta + dN_N^*) + 2d(\beta + dN_N^*)^2}. \tag{S15}$$

#### 1.3.2 Invasion of the non-infanticidal strategy into $E_I$

When a non-infanticidal mutant is rare in a resident population at the infanticidal boundary equilibrium $E_I$, the mutant variables are

$$\mathbf{y}_N = (C_{1,N}, C_{2,N}, M_N)^\top. \tag{S16}$$

The linearized mutant subsystem around $E_I$ is

$$\frac{d\mathbf{y}_N}{dt} = A_{N|I}\mathbf{y}_N, \tag{S17}$$

where

$$A_{N|I} = \begin{pmatrix} -(\beta + \gamma M_I^* + dN_I^*) & 0 & \dfrac{\alpha F_{f,I}^*}{M_I^*} \\ \beta & -(\beta + \gamma M_I^* + dN_I^*) & 0 \\ 0 & b_N\beta & -dN_I^* \end{pmatrix}. \tag{S18}$$

Thus, the invasion fitness of the non-infanticidal mutant is the eigenvalue of $A_{N|I}$ with the largest real part:

$$s_{N|I} = \max_j \operatorname{Re} \lambda_j \left(A_{N|I}\right). \tag{S19}$$

The characteristic polynomial of $A_{N|I}$ is

$$P_{N|I}(\lambda) = (\lambda + \beta + \gamma M_I^* + dN_I^*)^2(\lambda + dN_I^*) - b_N\beta^2\frac{\alpha F_{f,I}^*}{M_I^*}. \tag{S20}$$

For $b_N > 0$, $A_{N|I}$ is an irreducible Metzler matrix, and its dominant eigenvalue is real. Because $P_{N|I}(\lambda)$ is strictly increasing for $\lambda \geq 0$, the non-infanticidal strategy can invade if and only if $P_{N|I}(0) < 0$. Thus,

$$s_{N|I} > 0 \;\Leftrightarrow\; b_N\beta^2\frac{\alpha F_{f,I}^*}{M_I^*} > dN_I^*(\beta + \gamma M_I^* + dN_I^*)^2. \tag{S21}$$

Using the equilibrium relations at $E_I$, and $b_I = (1-s)b_N$, this condition can be written as

$$\frac{(\beta + dN_I^*)^2 + \frac{\gamma d}{2}(N_I^*)^2}{1-s} > \left(\beta + dN_I^* + \frac{\gamma N_I^*}{2}\right)^2. \tag{S22}$$

Rearranging gives

$$s > \frac{\frac{\gamma N_I^*}{2}\left(2\beta + dN_I^* + \frac{\gamma N_I^*}{2}\right)}{\left(\beta + dN_I^* + \frac{\gamma N_I^*}{2}\right)^2}. \tag{S23}$$

Note that $N_I^*$ depends implicitly on $s$ through the relationship $b_I = (1-s)b_N$.

### 2. two-species model

#### 2.1 non-evolutionary condition

To examine the ecological effect of male infanticide on population dynamics, we first consider a non-evolutionary scenario in which the male infanticidal strategy does not evolve. In this case, the frequency of infanticidal males in the male population is assumed to be constant.

Let $D_i = d_i(N_i + \rho_{ij}N_j)$ denote the density-dependent mortality rate of species $i$, where $j \neq i$. The population dynamics of species $i$ are then given by

$$\frac{dF_{f,i}}{dt} = -\alpha_i F_{f,i} + (1 + b_{N,i})\beta_i C_{2,N,i} + (1 + b_{I,i})\beta_i C_{2,I,i} - D_i F_{f,i}, \tag{S24a}$$

$$\frac{dC_{1,N,i}}{dt} = \alpha_i(1 - p_i)F_{f,i} - (\beta_i + \gamma_i p_i M_i)C_{1,N,i} - D_i C_{1,N,i}, \tag{S24b}$$

$$\frac{dC_{2,N,i}}{dt} = \beta_i C_{1,N,i} - (\beta_i + \gamma_i p_i M_i)C_{2,N,i} - D_i C_{2,N,i}, \tag{S24c}$$

$$\frac{dC_{1,I,i}}{dt} = \alpha_i p_i F_{f,i} + \gamma_i p_i M_i(C_{1,N,i} + C_{2,N,i} + C_{2,I,i}) - \beta_i C_{1,I,i} - D_i C_{1,I,i}, \tag{S24d}$$

$$\frac{dC_{2,I,i}}{dt} = \beta_i C_{1,I,i} - (\beta_i + \gamma_i p_i M_i)C_{2,I,i} - D_i C_{2,I,i}, \tag{S24e}$$

$$\frac{dM_i}{dt} = \beta_i(b_{N,i}C_{2,N,i} + b_{I,i}C_{2,I,i}) - D_i M_i. \tag{S24f}$$

Here, $p_i$ is the fixed frequency of infanticidal males in species $i$.

#### 2.2 quasi-steady-state approximation

To derive a reduced ecological system, we first consider the dynamics of the adult sex ratio. Let

$$F_i = F_{f,i} + C_{1,N,i} + C_{2,N,i} + C_{1,I,i} + C_{2,I,i} \tag{S25}$$

be the total female density of species $i$. Summing the equations for the five female states gives

$$\frac{dF_i}{dt} = \beta_i(b_{N,i}C_{2,N,i} + b_{I,i}C_{2,I,i}) - D_i F_i. \tag{S26}$$

Because male density satisfies

$$\frac{dM_i}{dt} = \beta_i(b_{N,i}C_{2,N,i} + b_{I,i}C_{2,I,i}) - D_i M_i, \tag{S27}$$

the difference between female and male densities follows

$$\frac{d}{dt}(F_i - M_i) = -D_i(F_i - M_i). \tag{S28}$$

Thus, when $(D_i > 0)$, deviations from an equal adult sex ratio decay over time. In the reduced ecological model, we therefore restrict the dynamics to the equal-sex-ratio,

$$F_i = M_i = \frac{N_i}{2}. \tag{S29}$$

We further assume that the distribution of females among reproductive states relaxes rapidly relative to changes in the total population density. We therefore apply the quasi-steady-state approximation to the fractions of females in each reproductive state. Define

$$x_{0,i} = \frac{F_{f,i}}{F_i}, x_{1,N,i} = \frac{C_{1,N,i}}{F_i}, x_{2,N,i} = \frac{C_{2,N,i}}{F_i}, x_{1,I,i} = \frac{C_{1,I,i}}{F_i}, x_{2,I,i} = \frac{C_{2,I,i}}{F_i}, \tag{S30}$$

which satisfy

$$x_{0,i} + x_{1,N,i} + x_{2,N,i} + x_{1I,i} + x_{2I,i} = 1. \tag{S31}$$

For a fixed frequency of infanticidal males, $p_i$, the infanticide rate experienced by a caring female is

$$h_i = \gamma_i p_i M_i = \frac{\gamma_i p_i N_i}{2}. \tag{S32}$$

The dynamics of the female-state fractions are then given by

$$\frac{dx_{0,i}}{dt} = -\alpha_i x_{0,i} + (1 + b_{N,i})\beta_i x_{2,N,i} + (1 + b_{I,i})\beta_i x_{2,I,i} - \beta_i(b_{N,i}x_{2,N,i} + b_{I,i}x_{2,I,i})x_{0,i}, \tag{S33a}$$

$$\frac{dx_{1,N,i}}{dt} = \alpha_i(1 - p_i)x_{0,i} - (\beta_i + h_i + \beta_i b_{N,i}x_{2,N,i} + \beta_i b_{I,i}x_{2,I,i})x_{1,N,i}, \tag{S33b}$$

$$\frac{dx_{2,N,i}}{dt} = \beta_i x_{1,N,i} - (\beta_i + h_i + \beta_i b_{N,i}x_{2,N,i} + \beta_i b_{I,i}x_{2,I,i})x_{2,N,i}, \tag{S33c}$$

$$\frac{dx_{1,I,i}}{dt} = \alpha_i p_i x_{0,i} + h_i(x_{1,N,i} + x_{2,N,i} + x_{2,I,i}) - \beta_i(b_{N,i}x_{2N,i} + b_{I,i}x_{2,I,i} + 1)x_{1,I,i}, \tag{S33d}$$

$$\frac{dx_{2,I,i}}{dt} = \beta_i x_{1,I,i} - (\beta_i + h_i + \beta_i b_{N,i}x_{2,N,i} + \beta_i b_{I,i}x_{2,I,i})x_{2,I,i}. \tag{S33e}$$

Under the quasi-steady-state approximation, these fractions are assumed to rapidly approach a stationary reproductive-state distribution. Thus, for each $N_i$ and fixed $p_i$, we determine

$$\boldsymbol{x}_i^* = (x_{0,i}^*, x_{1,N,i}^*, x_{2,N,i}^*, x_{1,I,i}^*, x_{2,I,i}^*) \tag{S34}$$

by solving

$$\frac{dx_{0,i}}{dt} = \frac{dx_{1,N,i}}{dt} = \frac{dx_{2,N,i}}{dt} = \frac{dx_{1,I,i}}{dt} = \frac{dx_{2,I,i}}{dt} = 0, \tag{S35}$$

together with

$$x_{0,i}^* + x_{1,N,i}^* + x_{2,N,i}^* + x_{1,I,i}^* + x_{2,I,i}^* = 1. \tag{S36}$$

The total population dynamics can then be approximated by

$$\frac{dN_i}{dt} = N_i[\beta_i(b_{N,i}x_{2,N,i}^* + b_{I,i}x_{2,I,i}^*) - d_i(N_i + \rho_{ij}N_j)]. \tag{S37}$$

Thus, for fixed $p_1$ and $p_2$, the original stage-structured system is reduced to a two-dimensional ecological system for $N_1$ and $N_2$, with reproductive-state structure and infanticide incorporated through the quasi-steady-state fractions $\boldsymbol{x}_i^*(N_i, p_i)$. We numerically solved Eqs. S35 and S36 for the stationary reproductive-state distribution. The resulting reduced equations were then evaluated to obtain the vector field and nullclines and to construct the phase-plane diagrams.

We further consider a simplified case, $s = 0$. Under this assumption, fecundity is independent of male strategy, $b_i = b_{N,i} = b_{I,i}$, allowing the female care classes to be aggregated across paternal types:

$$C_{1,i} = C_{1,N,i} + C_{1,I,i}, C_{2,i} = C_{2,N,i} + C_{2,I,i}. \tag{S38}$$

Writing

$$z_i^* = x_{2,i}^* = x_{2,N,i}^* + x_{2,I,i}^*, \tag{S39}$$

the quasi-steady-state fraction of females in the second care stage is given by the biologically feasible root of

$$b_i^2\beta_i^2(z_i^*)^3 + b_i\beta_i(\alpha_i + 2\beta_i + h_i)(z_i^*)^2 + (\beta_i^2 + 2\alpha_i\beta_i + \alpha_i h_i)z_i^* - \alpha_i\beta_i = 0. \tag{S40}$$

For $\alpha_i > 0$, $\beta_i > 0$, and $b_i > 0$, the left-hand side is strictly increasing for $z_i^* \geq 0$, and therefore the equation has a unique positive root. The reduced population dynamics are then

$$\frac{dN_i}{dt} = N_i[b_i\beta_i z_i^*(N_i, p_i) - d_i(N_i + \rho_{ij}N_j)]. \quad \text{(S41)}$$

Thus, in this special case, the effects of reproductive-state structure and infanticide on population growth are summarized by the single quasi-steady-state variable $z_i^*(N_i, p_i)$, representing the fraction of females in the second care stage.

**2.3 Mutual invasibility analysis with fixed strategy frequencies**

When infanticide occurs through density-dependent encounters, its demographic cost is weak when a species is rare but becomes progressively stronger as the species becomes abundant. Infanticide can therefore strengthen intraspecific self-limitation relative to interspecific competition, thereby generating a rare-species advantage. To isolate this effect from the direct fecundity cost of the infanticidal strategy, we consider the simplified case $s = 0$, so that $b_{N,i} = b_{I,i} = b_i$.

Let

$$\lambda_i(N_i, p_i) = b_i\beta_i z_i^*(N_i, p_i) \quad \text{(S42)}$$

denote the effective per-capita recruitment rate of species $i$, where $z_i^*$ is the quasi-steady-state fraction of females in the second care stage. The reduced population dynamics are

$$\frac{dN_i}{dt} = N_i[\lambda_i(N_i, p_i) - d_i(N_i + \rho_{ij}N_j)]. \quad \text{(S43)}$$

Holding the frequency of infanticidal males $p_i$ fixed, the infanticide hazard experienced by caring females increases with conspecific density. A higher hazard causes more frequent interruption of ongoing care and increases the expected time required to complete reproduction. Consequently, the fraction of females reaching the second care stage decreases with $N_i$, implying

$$\frac{\partial\lambda_i}{\partial N_i} = b_i\beta_i\frac{\partial z_i^*}{\partial N_i} < 0 \text{ for } p_i > 0. \quad \text{(S44)}$$

Thus, the effect of conspecific density on per-capita population growth is

$$\frac{\partial}{\partial N_i}\left(\frac{1}{N_i}\frac{dN_i}{dt}\right) = \frac{\partial \lambda_i}{\partial N_i} - d_i, \tag{S45}$$

whereas the corresponding effect of heterospecific density is

$$\frac{\partial}{\partial N_j}\left(\frac{1}{N_i}\frac{dN_i}{dt}\right) = -d_i \rho_{ij}. \tag{S46}$$

When species $i$ is rare, infanticidal encounters become negligible and this additional demographic cost largely disappears; when it is common, the cost becomes stronger. This asymmetry can generate a rare-species advantage and thereby broaden the conditions under which mutual invasibility and stable coexistence are possible [6,7].

When species $i$ is rare, the rate of infanticidal encounters approaches zero. Consequently, its low-density recruitment rate, denoted by $\lambda_i^0$, is independent of the frequency of infanticidal males within species $i$. The invasion growth rate of species $i$ into a resident population of species $j$ is therefore

$$r_i^{\text{inv}}(p_j) = \frac{1}{N_i}\frac{dN_i}{dt}\bigg|_{N_i \to 0,\ N_j = \hat{N}_j(p_j)} = \lambda_i^0 - d_i \rho_{ij} \hat{N}_j(p_j), \tag{S47}$$

where $\hat{N}_j(p_j)$ is the single-species equilibrium density of resident species $j$.

An increase in the frequency of infanticidal males in the resident species, $p_j$, reduces its equilibrium density, such that $d\hat{N}_j/dp_j < 0$. It follows that

$$\frac{\partial r_i^{\text{inv}}}{\partial p_j} = -d_i \rho_{ij} \frac{d\hat{N}_j}{dp_j} > 0. \tag{S48}$$

Thus, a higher frequency of infanticidal males in the resident species increases the invasion growth rate of its competitor.

For example, suppose that, in the absence of infanticide, species 1 is competitively superior to species 2. Species 1 can then invade the single-species equilibrium of species 2, whereas species 2 cannot invade that of species 1:

$$r_1^{\text{inv}}(0) = \lambda_1^0 - d_1 \rho_{12} \hat{N}_2(0) > 0, \tag{S49a}$$

$$r_2^{\text{inv}}(0) = \lambda_2^0 - d_2 \rho_{21} \hat{N}_1(0) < 0. \tag{S49b}$$

However, if an increase in $p_1$ sufficiently reduces the equilibrium density of species 1, the invasion growth rate of species 2 becomes positive, whereas that of species 1 does not change:

$$r_1^{\mathrm{inv}}(0) = \lambda_1^0 - d_1 \rho_{12} \hat{N}_2(0) > 0, \tag{S50a}$$

$$r_2^{\mathrm{inv}}(p_1) = \lambda_2^0 - d_2 \rho_{21} \hat{N}_1(p_1) > 0. \tag{S50b}$$

Species 2 can then invade the resident population of species 1. If species 1 also retains a positive invasion growth rate into the single-species equilibrium of species 2, the two species are mutually invasible. Density-dependent infanticide can therefore promote mutual invasibility by reducing the equilibrium density of the resident species and thereby releasing its competitor from interspecific competition. In this sense, infanticide strengthens the rare-species advantage associated with negative frequency dependence and broadens the conditions under which stable coexistence can occur.

### 3. Alternative forms of density dependence

#### 3.1 Density-independent mating initiation and infanticide

In this model, free females initiate reproduction at the density-independent rate $\alpha_i$. The paternal type of a newly initiated brood is assigned according to the current frequencies of the two male strategies. Thus, a proportion $1 - p_i$ of newly initiated broods is sired by non-infanticidal males, whereas a proportion $p_i$ is sired by infanticidal males. Infanticide is also independent of male density. Each caring female experiences an infanticide rate $\gamma_i p_i$, where $\gamma_i$ is the hazard when all males are infanticidal.

The eco-evolutionary dynamics of species $i$ are written as

$$\frac{d\boldsymbol{x}_i}{dt} = \boldsymbol{f}_i(\boldsymbol{x}_i, \boldsymbol{x}_j) = (f_{1,i}, f_{2,i}, f_{3,i}, f_{4,i}, f_{5,i}, f_{6,i}, f_{7,i})^{\top}, \tag{S51}$$

where

$$\frac{dF_{f,i}}{dt} = -\alpha_i F_{f,i} + (1 + b_{N,i})\beta_i C_{2,N,i} + (1 + b_{I,i})\beta_i C_{2,I,i} - D_i F_{f,i} \equiv f_{1,i}, \tag{S52a}$$

$$\frac{dC_{1,N,i}}{dt} = \alpha_i (1 - p_i) F_{f,i} - \beta_i C_{1,N,i} - \gamma_i p_i C_{1,N,i} - D_i C_{1,N,i} \equiv f_{2,i}, \tag{S52b}$$

$$\frac{dC_{2,N,i}}{dt} = \beta_i C_{1,N,i} - \beta_i C_{2,N,i} - \gamma_i p_i C_{2,N,i} - D_i C_{2,N,i} \equiv f_{3,i}, \tag{S52c}$$

$$\frac{dC_{1,I,i}}{dt} = \alpha_i p_i F_{f,i} + \gamma_i p_i (C_{1,N,i} + C_{2,N,i} + C_{2,I,i}) - \beta_i C_{1,I,i} - D_i C_{1,I,i} \equiv f_{4,i}, \tag{S52d}$$

$$\frac{dC_{2,I,i}}{dt} = \beta_i C_{1,I,i} - \beta_i C_{2,I,i} - \gamma_i p_i C_{2,I,i} - D_i C_{2,I,i} \equiv f_{5,i}, \tag{S52e}$$

$$\frac{dM_{N,i}}{dt} = b_{N,i} \beta_i C_{2,N,i} - D_i M_{N,i} \equiv f_{6,i}, \tag{S52f}$$

$$\frac{dM_{I,i}}{dt} = b_{I,i} \beta_i C_{2,I,i} - D_i M_{I,i} \equiv f_{7,i}. \tag{S52g}$$

Following the procedure described in Section S2.2, we derived a reduced model using a quasi-steady-state approximation based on a separation of timescales. We used the reduced model to calculate nullclines, boundary and interior equilibria, and invasion growth rates, and to construct phase-plane diagrams.

In the reduced model with fixed male-strategy frequencies, no stable coexistence equilibrium was found over the parameter range examined (Fig. S3).

In the full eco-evolutionary model, the evolution of infanticide slightly expanded the parameter region supporting stable coexistence beyond the coexistence boundary of the no-infanticide reference model (red region in Fig. S4A). In this additional coexistence region, bistability occurred between a locally stable coexistence equilibrium and a locally stable competitive-exclusion equilibrium. We also identified parameter regions with multiple locally stable coexistence equilibria, with up to four such equilibria (blue regions in Fig. S4A). The evolutionary outcomes at stable coexistence equilibria varied with the fecundity cost of the infanticidal strategy and the interspecific competition coefficient. However, no stable polymorphisms were observed, and only fixation of either the infanticidal or non-infanticidal strategy occurred (Fig. S4B). in another parameter region that partly overlapped with the baseline coexistence region, multiple equilibria other than the extinction equilibrium existed, but none were locally stable (purple region in Fig. S4A).

Overall, these results indicate that density-independent infanticide can alter the existence and stability of species coexistence equilibria but, unlike in the main model, does not consistently promote species coexistence through density-dependent self-limitation.

#### 3.2 Density-dependent mating initiation and infanticide

In this model, both mating initiation and infanticide arise through mass-action encounters between females and males. A free female initiates reproduction at the density-dependent per-capita rate $\alpha_i M_i$. Because encounters with the two male strategies occur in proportion to their respective densities, the flow of free females into broods sired by non-infanticidal males is $\alpha_i M_{N,i} F_{f,i}$, whereas the corresponding flow into broods sired by infanticidal males is $\alpha_i M_{I,i} F_{f,i}$. Infanticide is also assumed to be density-dependent. Each caring female encounters infanticidal males according to mass-action kinetics and therefore the total rate of infanticide in a female care class $C$ is $\gamma_i M_{I,i} C$. Thus, both reproductive initiation and infanticide become more frequent as the relevant male densities increase.

The eco-evolutionary dynamics of species $i$ are written as

$$\frac{d\boldsymbol{x}_i}{dt} = \boldsymbol{f}_i(\boldsymbol{x}_i, \boldsymbol{x}_j) = (f_{1,i}, f_{2,i}, f_{3,i}, f_{4,i}, f_{5,i}, f_{6,i}, f_{7,i})^\top, \tag{S53}$$

where each component is given by

$$\frac{dF_{f,i}}{dt} = -\alpha_i (M_{N,i} + M_{I,i}) F_{f,i} + (1 + b_{N,i}) \beta_i C_{2,N,i} + (1 + b_{I,i}) \beta_i C_{2,I,i} - D_i F_{f,i} \equiv f_{1,i} \tag{S54a}$$

$$\frac{dC_{1,N,i}}{dt} = \alpha_i M_{N,i} F_{f,i} - \beta_i C_{1,N,i} - \gamma_i M_{I,i} C_{1,N,i} - D_i C_{1,N,i} \equiv f_{2,i} \tag{S54b}$$

$$\frac{dC_{2,N,i}}{dt} = \beta_i C_{1,N,i} - \beta_i C_{2,N,i} - \gamma_i M_{I,i} C_{2,N,i} - D_i C_{2,N,i} \equiv f_{3,i} \tag{S54c}$$

$$\frac{dC_{1,I,i}}{dt} = \alpha_i M_{I,i} F_{f,i} + \gamma_i M_{I,i} (C_{1,N,i} + C_{2,N,i} + C_{2,I,i}) - \beta_i C_{1,I,i} - D_i C_{1,I,i} \equiv f_{4,i} \tag{S54d}$$

$$\frac{dC_{2,I,i}}{dt} = \beta_i C_{1,I,i} - \beta_i C_{2,I,i} - \gamma_i M_{I,i} C_{2,I,i} - D_i C_{2,I,i} \equiv f_{5,i} \tag{S54e}$$

$$\frac{dM_{N,i}}{dt} = b_{N,i} \beta_i C_{2,N,i} - D_i M_{N,i} \equiv f_{6,i} \tag{S54f}$$

$$\frac{dM_{I,i}}{dt} = b_{I,i} \beta_i C_{2,I,i} - D_i M_{I,i} \equiv f_{7,i} \tag{S54g}$$

Following the procedure described in Section S2.2, we derived a reduced model using a quasi-steady-state approximation based on a separation of timescales. We used the reduced model to calculate nullclines, boundary and interior equilibria, and invasion growth rates, and to construct phase-plane diagrams.

In the reduced model with fixed male-strategy frequencies, a locally stable interior coexistence equilibrium occurred over part of the parameter space examined (Fig. S5). However, this equilibrium coexisted with two locally stable species-exclusion equilibria, resulting in multistability. This is because the per-capita mating-initiation rate approaches zero as a species becomes rare, whereas mortality due to interspecific competition remains positive. Consequently, a species introduced at sufficiently low density cannot invade an established resident population, giving rise to priority effects [8] .

In the full eco-evolutionary model, the evolution of infanticide expanded the parameter region supporting locally stable coexistence beyond the coexistence boundary of the no-infanticide reference model (purple region in Fig. S6A). In this additional coexistence region, a locally stable coexistence equilibrium coexisted with two locally stable species-exclusion equilibria. More generally, both species-exclusion equilibria remained locally asymptotically stable throughout the parameter range examined, and the system exhibited up to three locally stable equilibria (Fig. S6A). The evolutionary outcomes at the stable coexistence equilibria varied with the fecundity cost and interspecific competition coefficient, ranging from fixation of the infanticidal strategy in both species to stable polymorphisms or fixation of the non-infanticidal strategy in one or both species (Fig. S6B).

Overall, these results indicate that, when both mating initiation and infanticide are density dependent, the evolution of infanticide can expand the parameter region supporting locally stable species coexistence. However, unlike in the main model, coexistence is not protected by mutual invasibility and remains dependent on initial conditions because the species-exclusion equilibria are also locally stable.

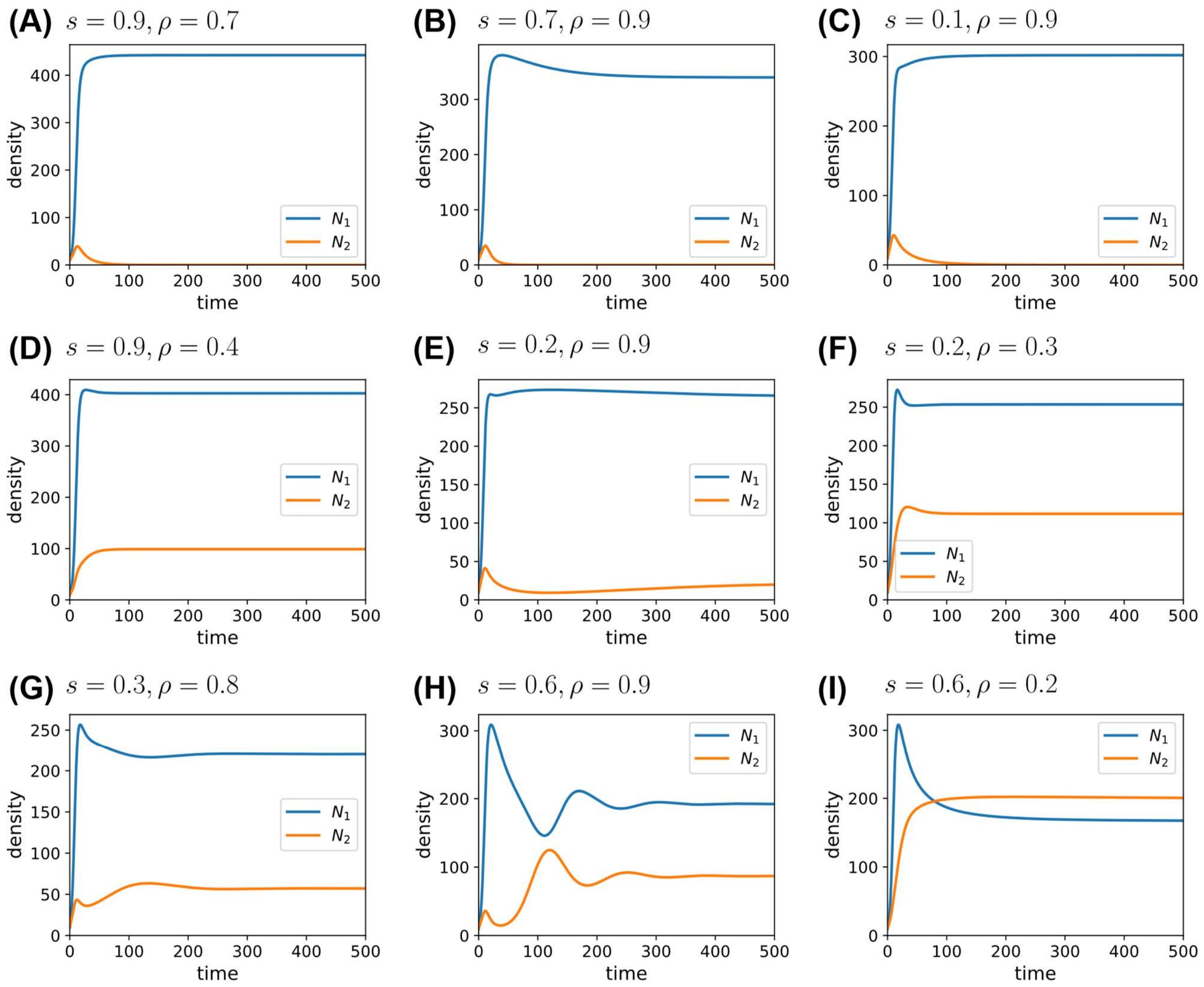


**Figure S1. Population dynamics of the full two-species eco-evolutionary model across nine parameter combinations.**

Panels (A–I) show the total adult densities of species 1 and species 2, $N_1$ and $N_2$, respectively. The infanticide cost $s = s_1 = s_2$ and competition coefficient $\rho = \rho_{12} = \rho_{21}$ are: (A) $s = 0.9, \rho = 0.7$; (B) $s = 0.7, \rho = 0.9$; (C) $s = 0.1, \rho = 0.9$; (D) $s = 0.9, \rho = 0.4$; (E) $s = 0.2, \rho = 0.9$; (F) $s = 0.2, \rho = 0.3$; (G) $s = 0.3, \rho = 0.8$; (H) $s = 0.6, \rho = 0.9$; and (I) $s = 0.6, \rho = 0.2$. Species 2 is competitively excluded in (A–C), whereas both species coexist in (D–I). Corresponding evolutionary trajectories are shown in Figure S2. All simulations start from $N_1 = N_2 = 10$ and $p_1 = p_2 = 0.5$, with equal total male and female densities and equal densities across the five female states. Other parameters are $b_{N,1} = 2.0$, $b_{N,2} = 1.0$, $\alpha_1 = \alpha_2 = 1$, $\beta_1 = \beta_2 = 1$, $\gamma_1 = \gamma_2 = 0.01$, and $d_1 = d_2 = 0.001$.

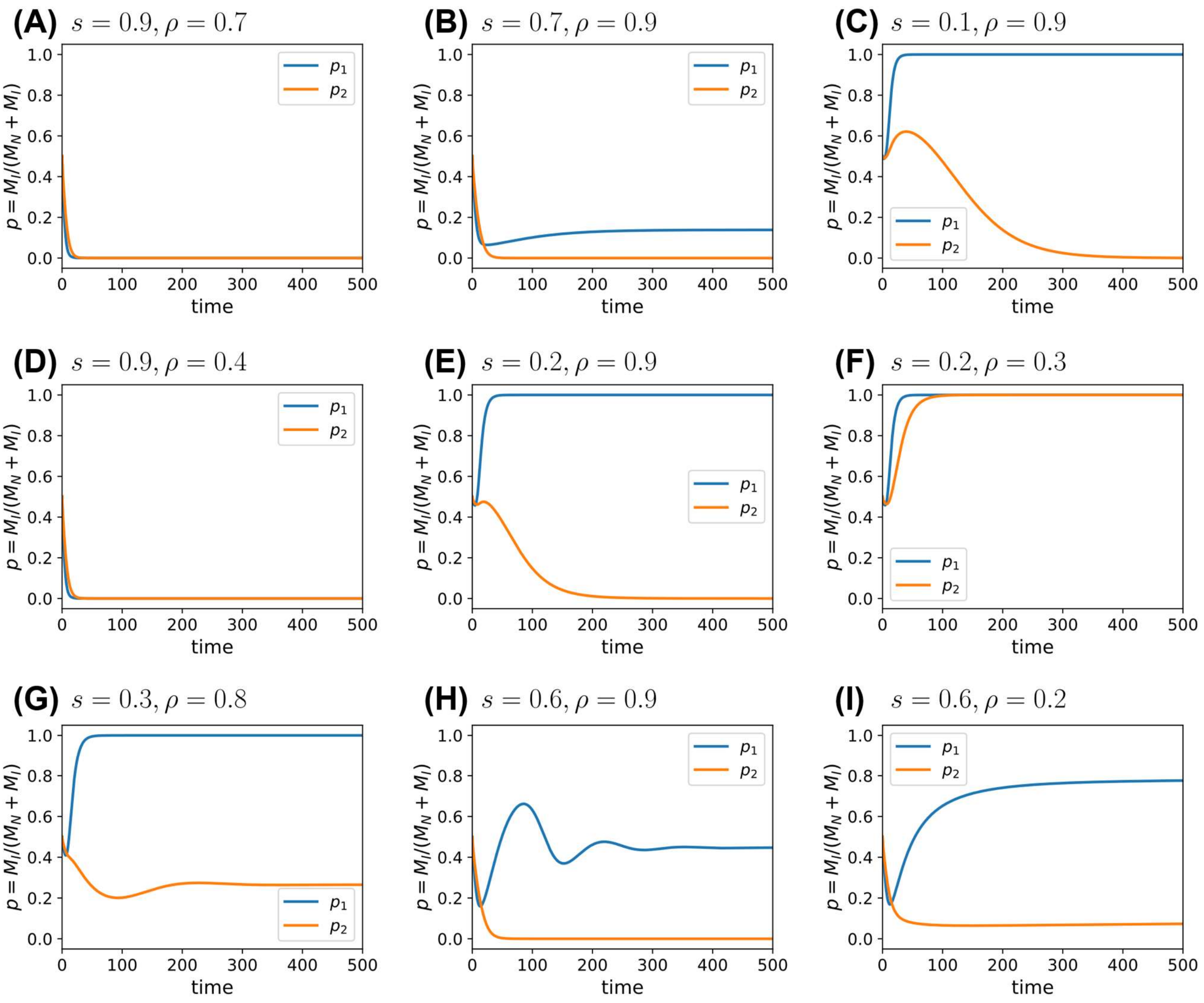


**Figure S2. Evolutionary dynamics of the full two-species eco-evolutionary model across nine parameter combinations.**

Panels (A–I) show the frequencies of infanticidal males in species 1 and species 2, $p_1$ and $p_2$, respectively. Panel ordering, parameter values, and initial conditions match Figure S1. In the remaining species 1 following the exclusion of species 2, (A) the non-infanticidal strategy becomes fixed, (B) a stable polymorphism is maintained, and (C) the infanticidal strategy becomes fixed. In the coexistence cases, (D) the non-infanticidal strategy becomes fixed in both species; (E) the infanticidal strategy becomes fixed in species 1 and the non-infanticidal strategy in species 2; (F) the infanticidal strategy becomes fixed in both species; (G) species 1 approaches fixation of the infanticidal strategy while species 2 approaches a stable polymorphism; (H) species 1 approaches a stable polymorphism while the non-infanticidal strategy becomes fixed in species 2; and (I) both species approach stable polymorphisms. In (A–C), species 2 becomes extinct, so its infanticidal-male frequency is undefined at the extinction equilibrium. The plotted $p_2$ trajectories represent frequencies during population decline while male density remains positive, rather than an equilibrium frequency in a surviving population.

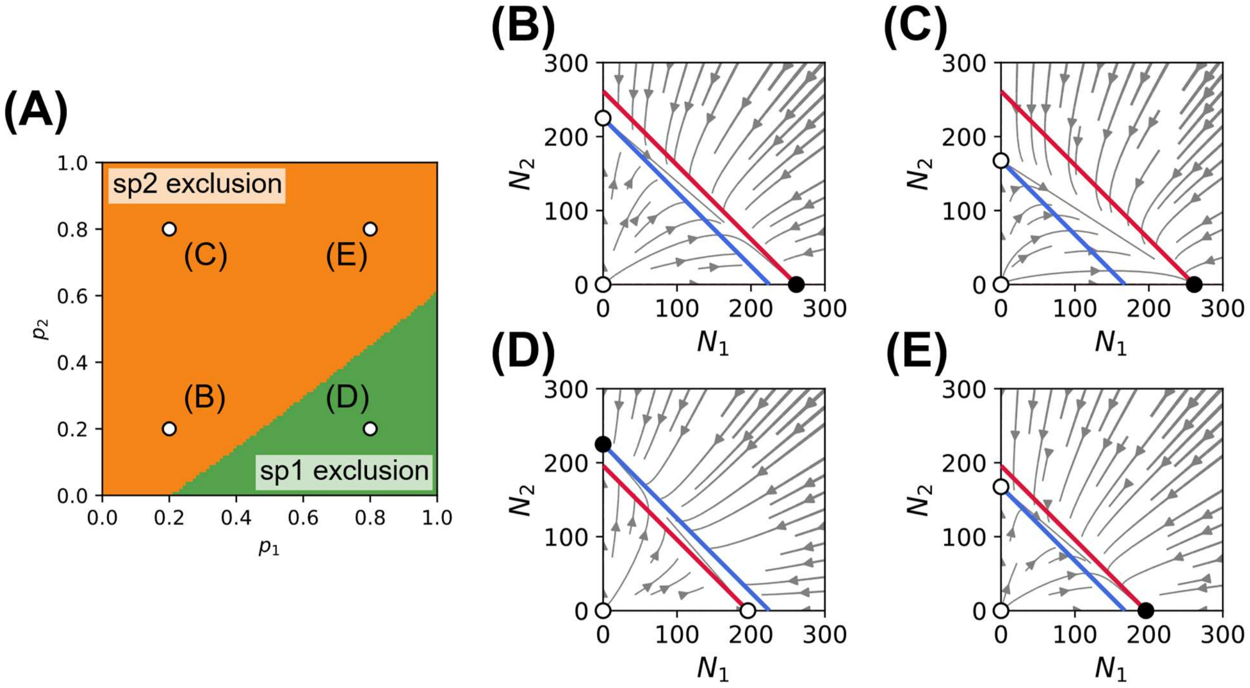


**Figure S3. Ecological outcomes and phase portraits of the reduced two-species model with density-independent mating and infanticide.**

Population dynamics were reduced to a two-dimensional system for the total population densities $N_1$ and $N_2$ using a time-scale separation approximation. In this model, both the transition of free females from $F_i$ to the first care stage $C_{1,i}$ and the occurrence of infanticide are density independent. The frequencies of infanticidal males, $p_1$ and $p_2$, were fixed and did not evolve. (A) Ecological outcomes as functions of $p_1$ and $p_2$. Green indicates competitive exclusion of species 1, and orange indicates competitive exclusion of species 2. Open circles in (A) indicate the parameter combinations illustrated in the phase portraits: (B) $(p_1, p_2) = (0.2, 0.2)$, (C) $(p_1, p_2) = (0.2, 0.8)$, (D) $(p_1, p_2) = (0.8, 0.2)$, and (E) $(p_1, p_2) = (0.8, 0.8)$. In panels (B–E), red and blue curves denote the non-trivial nullclines $dN_1/dt = 0$ and $dN_2/dt = 0$, respectively. Filled circles indicate locally stable equilibria, whereas open circles indicate unstable equilibria. Other parameters are $b_{N,1} = 1.2$, $b_{N,2} = 1.0$, $\alpha_1 = \alpha_2 = 1$, $s = 0.1$, $\beta_1 = \beta_2 = 1$, $\gamma_1 = \gamma_2 = 2$, $d_1 = d_2 = 0.001$, and $\rho_{12} = \rho_{21} = 1$.

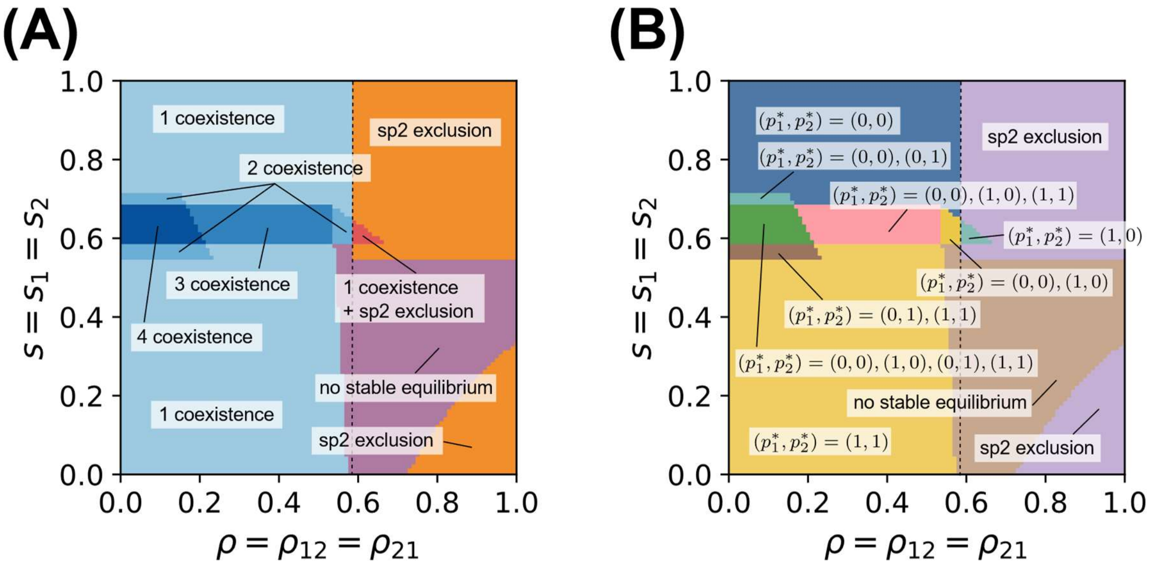


**Figure S4. Ecological and evolutionary outcomes of the full two-species model with density-independent mating and infanticide.**

Classification of locally asymptotically stable equilibria in the parameter plane defined by the interspecific competition coefficient, $\rho = \rho_{12} = \rho_{21}$, and the fecundity cost of the infanticidal strategy, $s = s_1 = s_2$. (A) Colors indicate the number and types of stable equilibria. When the infanticidal strategy is absent from both species, a unique stable coexistence equilibrium exists for $\rho \leq 0.58$ (dashed line). The evolution of infanticide altered the existence and stability of species coexistence equilibria but did not consistently promote species coexistence. (B) Colors indicate the equilibrium frequencies of infanticidal males in the coexistence equilibria. "No stable equilibrium" indicates that no locally asymptotically stable equilibrium was identified. Other parameters are $b_{N,1} = 2.0$, $b_{N,2} = 1.0$, $\alpha_1 = \alpha_2 = 1$, $\beta_1 = \beta_2 = 1$, $\gamma_1 = \gamma_2 = 1$, and $d_1 = d_2 = 0.001$.

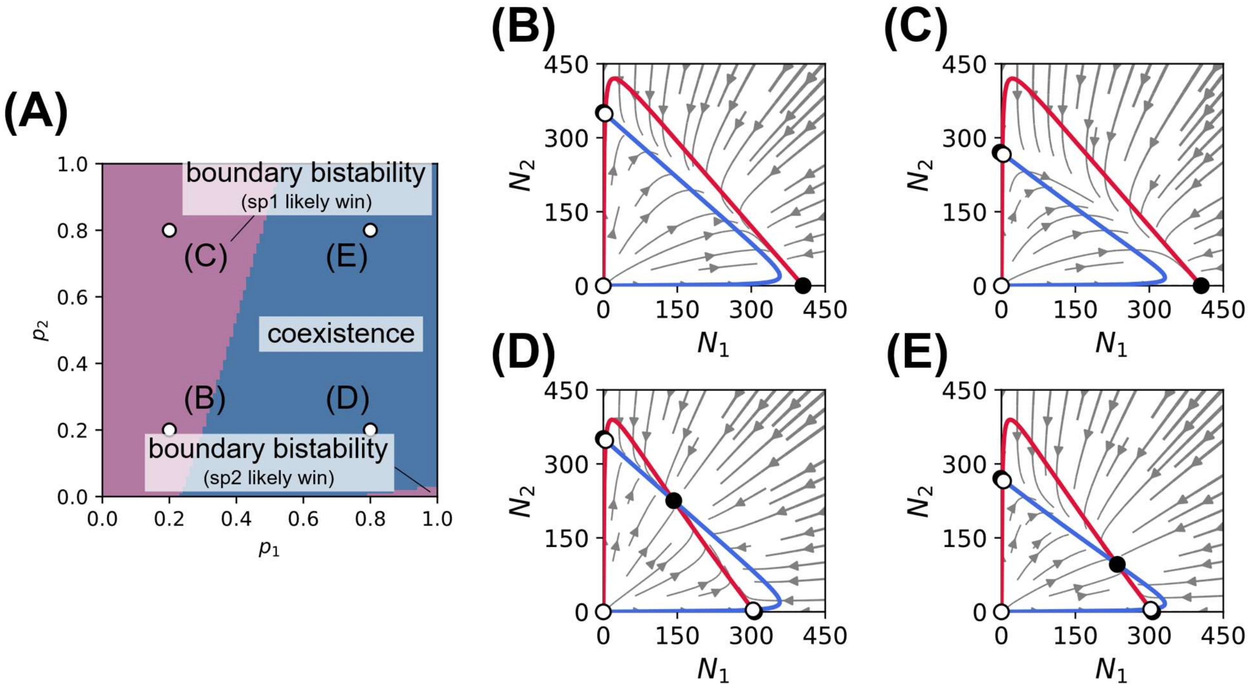


**Figure S5. Ecological outcomes and phase portraits of the reduced two-species model with density-dependent mating and infanticide.**

Population dynamics were reduced to a two-dimensional system for the total population densities $N_1$ and $N_2$ using a time-scale separation approximation. In this model, both the transition of free females from $F_i$ to the first care stage $C_{1,i}$ and the occurrence of infanticide are density dependent. The frequencies of infanticidal males, $p_1$ and $p_2$, were fixed and did not evolve. (A) Ecological outcomes as functions of $p_1$ and $p_2$. Blue indicates the existence of a locally stable coexistence equilibrium in addition to the two locally stable boundary equilibria. Purple indicates boundary bistability. Open circles in (A) indicate the parameter combinations illustrated in the phase portraits: (B) $(p_1, p_2) = (0.2,0.2)$, (C) $(p_1, p_2) = (0.2,0.8)$, (D) $(p_1, p_2) = (0.8,0.2)$, and (E) $(p_1, p_2) = (0.8,0.8)$. In panels (B–E), red and blue curves denote the non-trivial nullclines $dN_1/dt = 0$ and $dN_2/dt = 0$, respectively. Filled circles indicate locally stable equilibria, whereas open circles indicate unstable equilibria. Other parameters are $b_{N,1} = 1.2$, $b_{N,2} = 1.0$, $\alpha_1 = \alpha_2 = 1$, $s = 0.1$, $\beta_1 = \beta_2 = 1$, $\gamma_1 = \gamma_2 = 0.01$, $d_1 = d_2 = 0.001$, and $\rho_{12} = \rho_{21} = 1$.

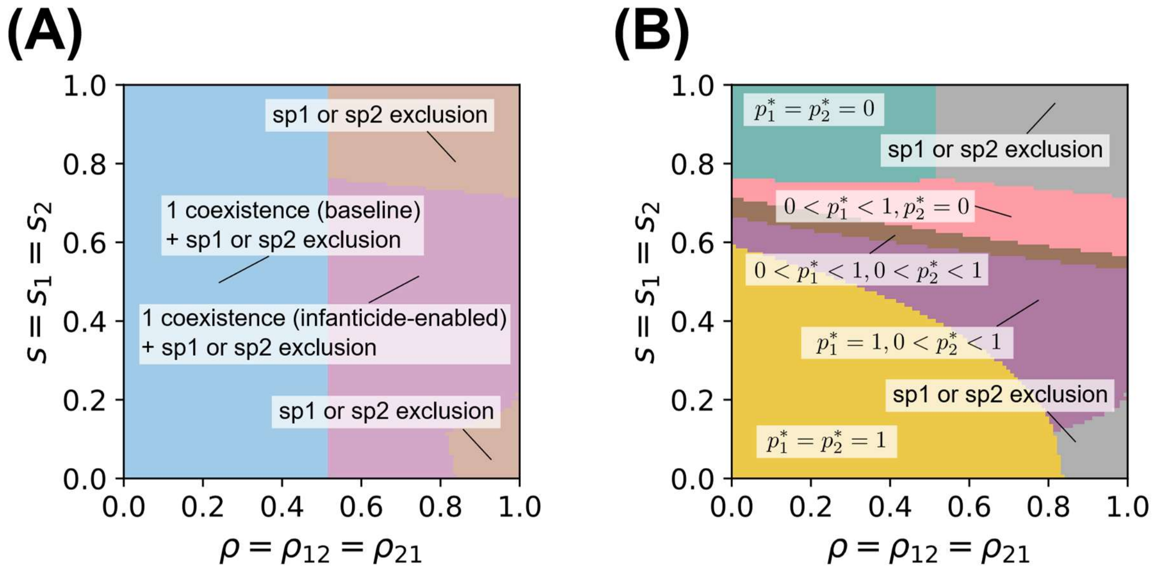


**Figure S6. Ecological and evolutionary outcomes in the full two-species model with density-dependent mating and infanticide.**

Classification of locally asymptotically stable equilibria in the parameter plane defined by the interspecific competition coefficient, $\rho = \rho_{12} = \rho_{21}$, and the fecundity cost of the infanticidal strategy, $s = s_1 = s_2$. (A) Colors indicate the number and types of stable equilibria. Both single-species equilibria were always locally asymptotically stable, and the system exhibited up to three locally asymptotically stable equilibria. Baseline coexistence refers to the region where a stable coexistence equilibrium exists even when both species consist entirely of non-infanticidal individuals. The evolution of infanticide expanded the parameter region in which a stable coexistence equilibrium exists (purple region). (B) Colors indicate the equilibrium frequencies of infanticidal males in the stable coexistence equilibria. Other parameters are $b_{N,1} = 2.0$, $b_{N,2} = 1.0$, $\alpha_1 = \alpha_2 = 1$, $\beta_1 = \beta_2 = 1$, $\gamma_1 = \gamma_2 = 0.01$, and $d_1 = d_2 = 0.001$.
